# Prefrontal Cortex as a Meta-Reinforcement Learning System

**DOI:** 10.1101/295964

**Authors:** Jane X. Wang, Zeb Kurth-Nelson, Dharshan Kumaran, Dhruva Tirumala, Hubert Soyer, Joel Z. Leibo, Demis Hassabis, Matthew Botvinick

## Abstract

Over the past twenty years, neuroscience research on reward-based learning has converged on a canonical model, under which the neurotransmitter dopamine ‘stamps in’ associations between situations, actions and rewards by modulating the strength of synaptic connections between neurons. However, a growing number of recent findings have placed this standard model under strain. In the present work, we draw on recent advances in artificial intelligence to introduce a new theory of reward-based learning. Here, the dopamine system trains another part of the brain, the prefrontal cortex, to operate as its own free-standing learning system. This new perspective accommodates the findings that motivated the standard model, but also deals gracefully with a wider range of observations, providing a fresh foundation for future research.

---

Exhilarating advances have recently been made toward understanding the mechanisms involved in reward-driven learning. This progress has been enabled in part by the importation of ideas from the field of reinforcement learning^1^ (RL). Most centrally, this input has led to an RL-based theory of dopaminergic function. Here, phasic dopamine (DA) release is interpreted as conveying a reward prediction error (RPE) signal^2–4^, an index of surprise which figures centrally in temporal-difference RL algorithms^1^. Under the theory, the RPE drives synaptic plasticity in the striatum, translating experienced action-reward associations into optimized behavioral policies^4, 5^. Over the past two decades, evidence has steadily mounted for this proposal, establishing it as the standard model of reward-driven learning.

However, even as this standard model has solidified, a collection of problematic observations has accumulated. One quandary arises from research on prefrontal cortex (PFC). A growing body of evidence suggests that PFC implements mechanisms for reward-based learning, performing computations that strikingly resemble those ascribed to DA-based RL. It has long been established that sectors of the PFC represent the expected values of actions, objects and states^6–8^. More recently, it has emerged that PFC also encodes the recent history of actions and rewards^9–15^. The set of variables encoded, along with observations concerning the temporal profile of neural activation in the PFC, has led to the conclusion that “PFC neurons dynamically [encode] conversions from reward and choice history to object value, and from object value to object choice”^10^. In short, neural activity in PFC appears to reflect a set of operations that together constitute a self-contained RL algorithm.

Placing PFC beside DA, we obtain a picture containing two full-fledged RL systems, one utilizing activity-based representations and the other synaptic learning. What is the relationship between these systems? If both support RL, are their functions simply redundant? One suggestion has been that DA and PFC subserve different forms of learning, with DA implementing *model-free* RL, based on direct stimulus-response associations, and PFC performing *model-based* RL, which leverages internal representations of task structure^16–18^. However, an apparent problem for this dual-system view is the repeated observation that DA prediction-error signals are informed by task structure, reflecting “inferred”^18, 19^ and “model-based”^20–22^ value estimates that are difficult to square with the standard theory as originally framed.

## A New Formulation

In the present work we offer a new perspective on the computations underlying reward-based learning, one that accommodates the findings that motivated the existing theories reviewed above, but which also resolves many of the prevailing quandaries. On a wider level, the theory suggests a coherent explanation for a diverse range of findings previously considered unconnected.

We begin with three key premises:

1. *System architecture:* In line with previous work^23–26^, we conceptualize the PFC, together with the basal ganglia and thalamic nuclei with which it connects, as forming a recurrent neural network. This network’s inputs include perceptual data, which either contains or is accompanied by information about executed actions and received rewards^11, 27, 28^. On the output side, the network triggers actions and, additionally, emits estimates of state value (Figure 1A,B).
2. *Learning:* As suggested in past research^29–31^, we assume that the synaptic weights within the prefrontal network, including its striatal components, are adjusted by a model-free RL procedure, within which DA conveys a RPE signal. Via this role, the DA-based RL procedure shapes the activation dynamics of the recurrent prefrontal network.
3. *Task environment:* Following past proposals^32–35^, we assume that RL takes place not on a single task, but instead in a dynamic environment posing a series of interrelated tasks. The learning system is thus required to engage in ongoing inference and behavioral adjustment.

**Figure 1.**
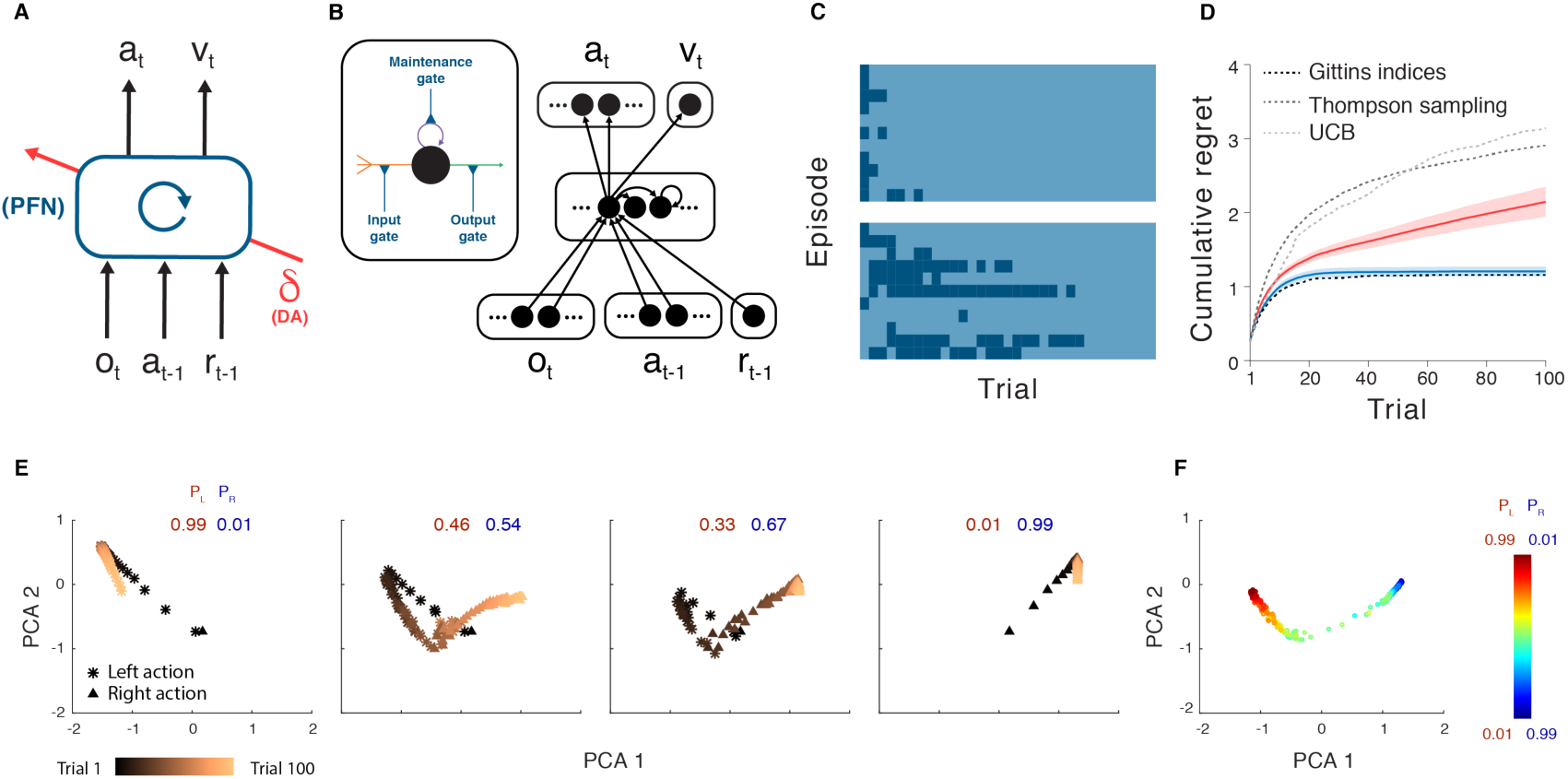
A. Agent architecture. The prefrontal network (PFN), including sectors of the basal ganglia and the thalamus that connect directly with PFC, is modeled as a recurrent neural network, with synaptic weights adjusted through an RL algorithm driven by DA. ***o*** = perceptual input, *a* = action, *r* = reward, *v* = state value, *t* = timestep, *δ* = RPE. The central box denotes a single, fully connected set of LSTM units. B. A more detailed schematic of the neural network implementation used in our simulations. Input units encoding the current observation, and previous action and reward connect all-to-all with hidden units, which are themselves fully connected LSTM units. These units connect all-to-all, in turn, with a softmax layer (see Methods) coding for actions, and a single linear unit coding for estimated state value. Inset: A single LSTM unit (see Hochreiter, et al^72^). Input (orange) is a weighted sum of other unit outputs, plus the activity of the LSTM unit itself (purple). Output (green) puts these summed inputs through a sigmoid non-linearity. All three quantities are multiplicatively gated (blue). Full details, including relevant equations are presented in Methods. C. Trial-by-trial model behavior on bandit problems with Bernoulli (arm 1, arm 2) reward parameters 0.25, 0.75 (top), 0.6, 0.4 (bottom). Contrasting colors demark left versus right actions. The network shifts from exploration to exploitation, making this transition more slowly in the more difficult problem. D. Performance for the meta-RL network trained on bandits with independently, identically distributed arm parameters, and tested on 0.25, 0.75 (red), measured in terms of cumulative regret, defined as the cumulative loss (in expected rewards) suffered when playing sub-optimal (lower-reward) arms. Performance of several standard machine-learning bandit algorithms is plotted for comparison. Blue: Performance on the same problem after training on correlated bandits (parameters always summed to 1). E. Evolution of RNN activation pattern during individual trials while testing on correlated bandits, after training on problems from the same distribution. *P_l_* = probability of reward for action *left*. *P_r_* = probability of reward for action *right*. F. RNN activity patterns from step 100 in the correlated bandit task, across a range of payoff parameters. Further analyses are presented in Supplementary Figure 1. Analysis done for 300 evaluation episodes, each consisting of 100 trials, performed by 1 fully trained network.

As indicated, these premises are all firmly grounded in existing research. The novel contribution of the present work is to identify an emergent effect that results when the three premises are concurrently satisfied. As we will show, these conditions, when they co-occur, are sufficient to produce a form of *meta-learning*^36^, where one learning algorithm gives rise to a second, more efficient learning algorithm. Specifically, by adjusting the connection weights within the prefrontal network, DA-based RL creates a second RL algorithm, implemented entirely in the prefrontal network’s activation dynamics. This new learning algorithm is independent of the original one, and differs in ways that are suited to the task environment. Crucially, the emergent algorithm is a full-fledged RL procedure: It copes with the exploration-exploitation tradeoff^37, 38^, maintains a representation of the value function^1^, and progressively adjusts the action policy. In view of this point, and in recognition of some precursor research^39–42^, we refer to the overall effect as *meta-reinforcement learning*.

## Meta-reinforcement learning: An illustrative example

For demonstration, we leverage the simple model shown in Figure 1A, a recurrent neural network (Figure 1B) whose weights are trained using a model-free RL algorithm (see Algorithm 1 and Methods), exploiting recent advances in deep learning research^43^. We consider this model’s performance in a simple ‘two-armed bandit’ RL task^41^. On each trial, the system outputs an action: *left* or *right*. Each has a probability of yielding a reward, but these probabilities change with each training episode, thus presenting a new bandit problem. After training on a series of problems, the weights in the recurrent network are fixed, and the system is tested on further problems. Figure 1C,D illustrates its performance. The network explores both arms, gradually honing in on the richer one, learning with an efficiency that rivals standard machine-learning algorithms.

Because the weights in the network were fixed at test, the system’s learning ability cannot be attributed to the RL algorithm that was used to tune the weights. Instead, learning reflects the activation dynamics of the recurrent network. As a result of training, these dynamics implement their own RL algorithm, integrating reward information over time, exploring, and refining the action policy (see Figure 1E,F and Supplementary Figure 1). This learned RL algorithm not only functions independently of the algorithm that was originally used to set the network weights; it also *differs* from that original algorithm in ways that make it specially adapted to the task distribution on which the system was trained. An illustration of this point is presented in Figure 1D, which shows performance of the same system after training on a structured version of the bandit problem, where the arm parameters were anti-correlated across episodes. Here, the recurrent network converges on an RL algorithm that exploits the problem’s structure, identifying the superior arm more rapidly than in the unstructured task.

## Neuroscientific Interpretation

Having introduced meta-RL in abstract computational terms, we now return to its neurobiological interpretation. This starts by regarding the prefrontal network, including its subcortical components, as a recurrent neural network. DA, as in the standard model, broadcasts an RPE signal, driving synaptic learning within the prefrontal network. The principal role of this learning is to shape the dynamics of the prefrontal network by tuning its recurrent connectivity. Through meta-RL, these dynamics come to implement a second RL algorithm, which differs from the original DA-driven algorithm, assuming a form tailored to the task environment. The role of DA-driven RL, under this account, plays out across extended series of tasks. Rapid within-task learning is mediated primarily by the emergent RL algorithm inherent in the dynamics of the prefrontal network.

## Results

Despite its simplicity, the meta-RL framework can account for a surprising range of neuroscientific findings, including many of the results that have presented difficulties for the standard RPE model of DA. To survey the framework’s ramifications we conducted a set of six simulation experiments, each focusing on a set of experimental findings chosen to illustrate a core aspect of the theory. Throughout these simulations, we continue to apply the simple computational architecture from Figure 1A. This approach obviously abstracts over many neuroanatomical and physiological details (see Supplementary Figure 8). However, it allows us to demonstrate key effects in a setting where their computational origins can be readily identified. In the same spirit, our simulations focus not on parameter-dependent fits to data, but instead only on robust qualitative effects.

### Simulation 1. Reinforcement learning in the prefrontal network

We begin with the finding that PFC encodes recent actions and rewards, integrating these into an evolving representation of choice value. To demonstrate how meta-RL explains this finding, we simulate results from Tsutsui and colleagues^10^ and Lau and Glimcher^44^. Both studies employed a task in which monkeys chose between two visual saccade targets, each of which yielded juice reward with a probability that intermittently changed. Given the reward schedule, an optimal strategy could be approximated by sampling each target in proportion to its yield^45^, and the monkeys displayed such ‘probability matching’ (Figure 2A). Recordings from dorsolateral PFC^10^ revealed activity encoding the target selected on the previous trial, the reward received, the updated values of the targets, and each upcoming action (Figure 2B), supporting the conclusion that PFC neurons reflect the trial-by-trial construction and updating of choice value from recent experience.^15^

**Figure 2.**
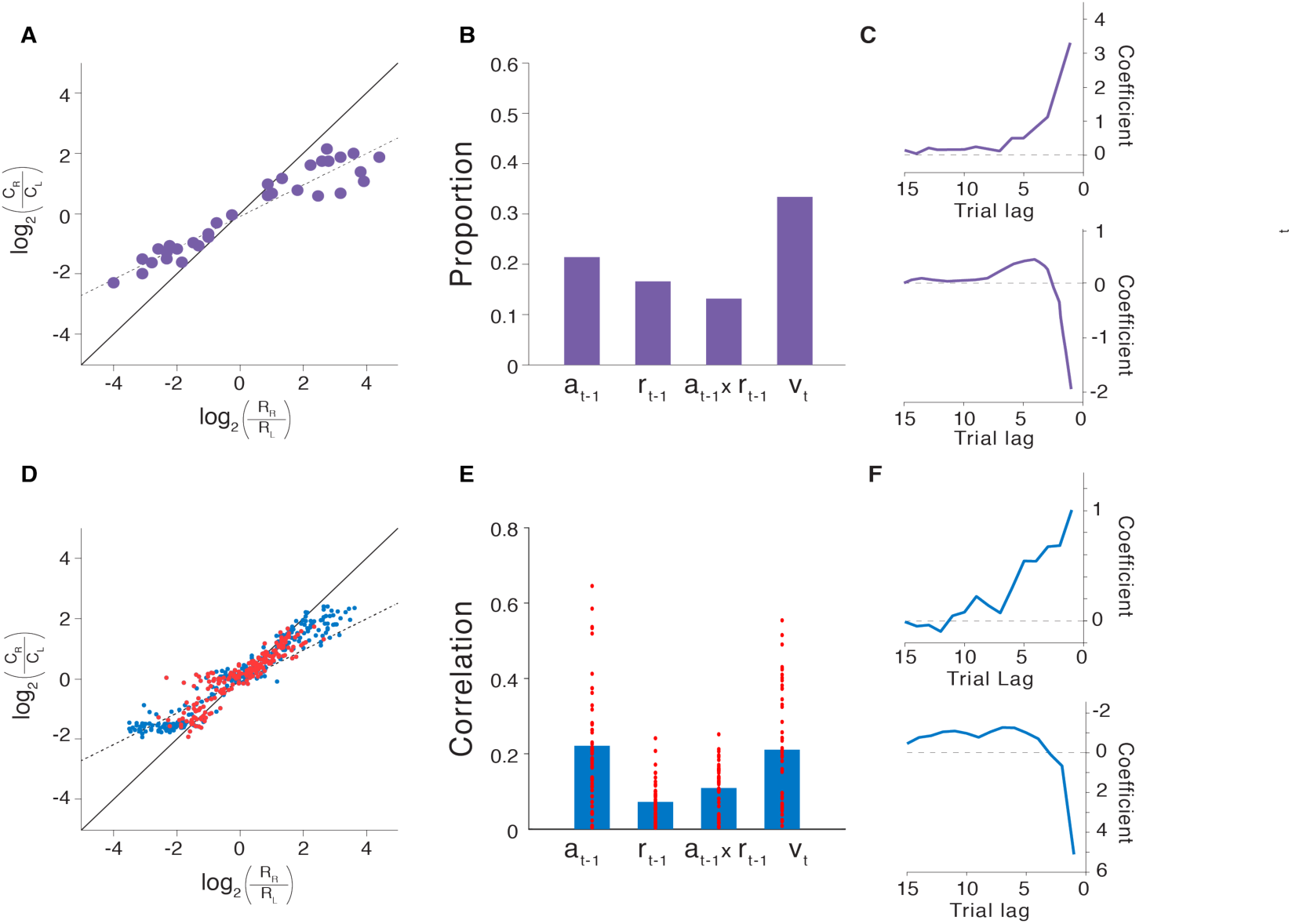
A. Probability matching behavior from Lau and Glimcher^44^. *C_i_*: Number of trials on which action *i* was selected. *R*: Number of reward events yielded by action *i*. B. Proportion of PFC neurons coding, during the trial-initial fixation period, for preceding action, preceding reward, the interaction of these two factors, and current choice value, from Tsutsui et al.^10^ C. Lag regression coefficients indicating the influence of recent reward outcomes (top) and actions (bottom) on choice from Lau and Glimcher^44^. D: Matching behavior from model. Red points show generalization performance (see Methods). E. Partial correlation coefficients (absolute values, Pearson’s r) for the same factors as those in panel B, but across units in the model at trial-initial fixation. Bars indicate mean values. Coding for the upcoming action (not shown) was also ubiquitous across units, and generally stronger than for the other variables. F. Regression weights corresponding to those in panels C, but based on model behavior. Differences in coefficient scale between panels C and F are due to minor differences in the regression procedure, specifically the use of binary indicators for reward history vs. continuous values (ranging from 0.35 to 0.6). For comparison, Tsutsui and colleagues reported regression results for the effect of past reward which indicted a similar temporal profile, but peaked at lag one with a coefficient of 1.25. Panels A-C adapted with permission from Tsutsui et al^10^. and Lau and Glimcher^44^.

We simulated these results by training our meta-RL model on the same task (see Methods). The network adapted its behavior to each task instantiation, displaying probability matching that closely paralleled experimental results (Figure 2D). Finer-grained analyses yielded patterns strikingly similar to those observed in monkey behavior (Figure 2C,F and Supplementary Figure 2). Note that the network’s weights were fixed at test, meaning that the network’s behavior can only result from the dynamics of prefrontal RNN activity. Consistent with this, we found units within the prefrontal network that coded for each of the variables reported by Tsutsui and colleagues (Figure 2E).

These results provide another basic illustration of meta-RL in action, with DA-driven training giving rise to an independent prefrontal-based learning algorithm. In this case, meta-RL accounts for both the behavior and PFC activity observed in the experimental data, but also offers an explanation for how these both may have emerged through DA-driven learning.

### Simulation 2. Adaptation of prefrontal-based learning to the task environment

A key aspect of meta-RL is that the learning algorithm that arises within the prefrontal network can differ from the DA-based algorithm that created it. We illustrate this principle here, focusing on differences in a single parameter: the learning rate. To this end, we simulate results from an experiment by Behrens and colleagues^46^. This involved a two-armed bandit task, which alternated between ‘stable’ periods where payoff probabilities held steady, and ‘volatile’ periods where they fluctuated. Behrens and colleagues found that human participants adopted a faster learning rate during volatile periods than stable periods, in accord with the optimal strategy identified by a Bayesian model which adjusted its learning rate based on dynamic estimates of task volatility (cf. Figure 3A,C). Neuroimaging data identified a region within PFC (specifically, dorsal cingulate cortex) whose activity tracked the Bayesian model’s volatility estimates.

**Figure 3.**
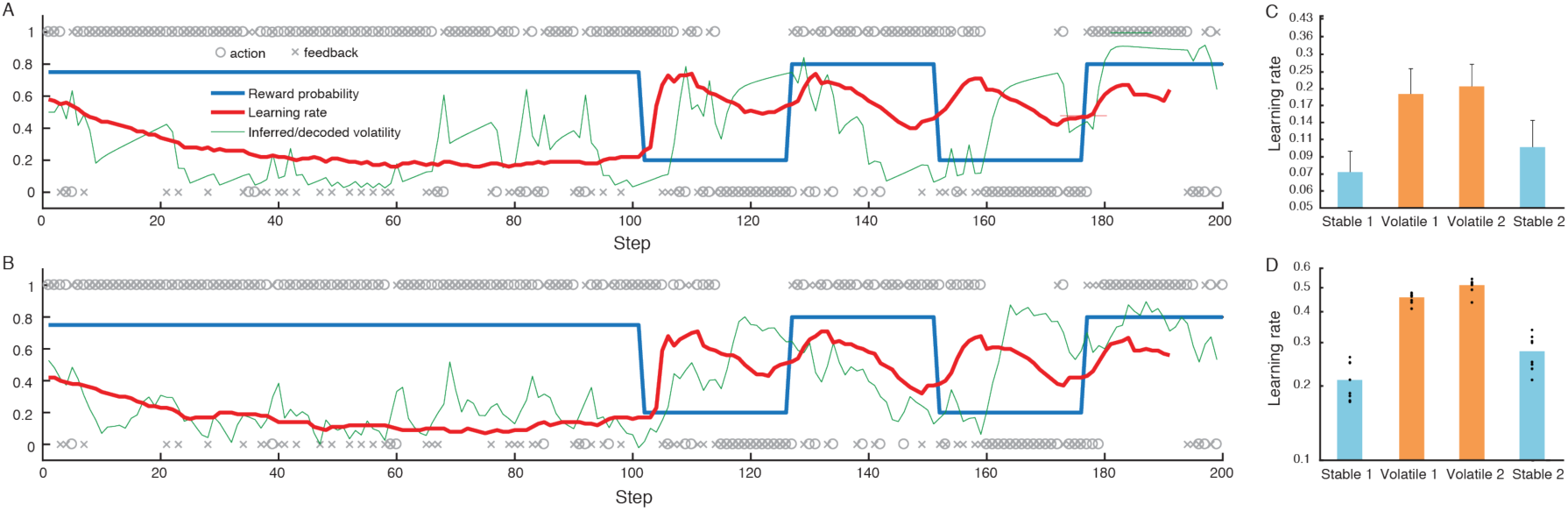
A. Sample behavior of the Bayesian model proposed by Behrens et al.^46^ on their volatile bandit task. Blue: True reward probability for action 1 (probabilities for actions 1 and 2 summed to 1). o = Actions (action 1 above, 2 below). x = Outcomes (reward above, non-reward below). Green: Estimated volatility. Red: Learning rate. B. Corresponding quantities for the meta-RL model, with volatility decoded as described in Methods. C. Summary of human behavior redrawn from Behrens et al.^46^, showing mean learning rates across test blocks with standard errors. D. Corresponding meta-RL behavior showing a similar pattern, with higher learning rates in volatile task blocks. Dots represent data from separate training runs (n = 8) employing different random seeds. Panels A and C adapted with permission from Behrens et al.^46^

Figure 3B,D summarizes the behavior of our model, with fixed weights, on the same task. The model was previously trained on a series of episodes with shifting volatilities (see Methods and Supplementary Figure 3) to mimic the prior experience of different volatilities that human participants brought to the task. Like human learners, the network dynamically adapted its learning rate to the changing volatility. Moreover, like the PFC activity observed with fMRI, many units within the LSTM (37±1%) explicitly tracked the changing volatility.

Critically, the learning rates indicated in Figure 3D are orders of magnitude larger than the one governing the DA-based RL algorithm that adjusted the prefrontal network’s weights during training (which was set at 0.00005). The results thus provide a concrete illustration of the point that the learning algorithm which arises from meta-RL can differ from the algorithm that originally engendered it. The results also allow us to emphasize an important corollary of this principle, which is that meta-RL produces a prefrontal learning algorithm that is *adapted* to the task environment. In the present case, this adaptation manifests in the way the learning rate responds to task fluctuations. Previous studies have proposed special-purpose mechanisms to explain dynamic shifts in learning rate^39, 47, 48^. Meta-RL explains these shifts as an emergent effect, arising from a very general set of conditions. Moreover, dynamic learning rates constitute just one possible form of specialization. When meta-RL occurs in environments with different structures, qualitatively different learning rules will emerge, a point that will be illustrated by subsequent simulations.

### Simulation 3. Reward prediction errors reflecting inferred value

As introduced earlier, one apparent challenge for the standard model is that the DA RPE signal reflects knowledge of task structure. An example is the “inferred-value” effect reported by Bromberg-Martin and colleagues.^18^ In each trial in their task, a visual target appeared on either the left or right side of a display, and the monkey was expected to saccade to that target. At any point in the experiment, either the left or right target yielded juice reward while the other target did not, and these role assignments reversed intermittently throughout the testing session. The key observation concerned DA signals following a reversal: Immediately after the monkey experienced a change in reward for one target, the DA response to the appearance of the *other* target changed dramatically, reflecting an inference that the value of that target had also changed (Figure 4A).

**Figure 4.**
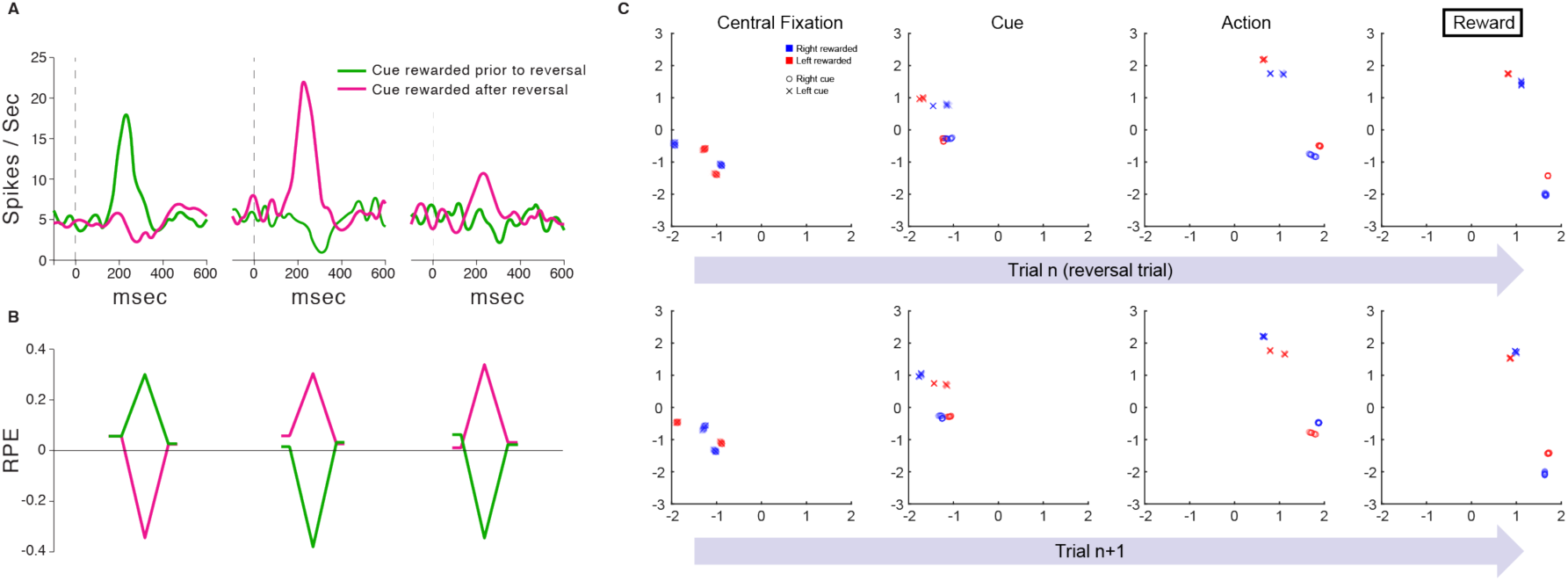
A. Results adapted with permission from Bromberg-Martin et al.^18^ Dopaminergic activity in response to cues ahead of a reversal (left) and for cues with an experienced (middle) and inferred (right) change in value. Curves represent average neuronal responses (n = 42 – 63 for each curve) from 2 monkeys. B. Corresponding RPE signals from the model. Leading and trailing points for each data-series correspond to initial fixation and saccade steps. Peaks/troughs correspond to stimulus presentation. Although the RPE responses in empirical data are smaller for inferred value, which is not shown here, other empirical results from Bromberg-Martin et al depict much more similar patterns. In particular, recordings from lateral habenula in the same task, but from different animals, yielded much more similar RPE signals (and behavioral reaction times) across conditions. C. First two principal components of RNN activity (LSTM output) on individual steps in the task from Simulation 3, analyzed from 1200 evaluation episodes performed by 1 trained network replica. Other replicas yielded very similar results. The top row focuses on reversal trials, where the final reward feedback (rightmost panel) signals that the rewarded target has switched, relative to preceding trials. The second row focuses on trials immediately following the reversal trials examined in the first row. Activity patterns cluster according to the current latent state of the task (i.e., which cue is currently rewarded), and later in each trial also according to the action selected. As shown in the leftmost panels (‘Central Fixation’), at the beginning of each trial, the network’s activation state clearly represents the latent state of the task, and this abruptly reverses following reversal trials.

This inferred-value effect, along with other related findings, has given rise to models in which either PFC or hippocampus encodes abstract latent-state representations^27, 49–54^, which can then feed into the computations generating the RPE^19^. As it turns out, a related mechanism arises naturally from meta-RL. To show this, we trained our meta-RL model on Bromberg-Martin’s task. At test, we observed RPE signals that reproduced the pattern displayed by DA. In particular, the model clearly reproduced the critical inferred-value effect (Figure 4B).

The explanation for this result is straightforward. In our architecture, the reward prediction component of the RPE comes from the prefrontal network’s state-value output, consistent with data showing that DA signaling is influenced by projections from PFC^19^. As DA-based training causes the prefrontal network to encode information about task dynamics (see Figure 4C), this information also manifests in the reward prediction component of the RPE.

### Simulation 4. ‘Model-based’ behavior: The Two-Step Task

Another important setting where structure-sensitive DA signaling has been observed is in tasks designed to probe for model-based control. Perhaps the most heavily studied task in this category is the ‘two-step’ paradigm introduced by Daw and colleagues^21^. In the version we will consider (Figure 5A), every trial begins with a decision between two actions. Each triggers a probabilistic transition to one of two perceptually distinguishable second-stage states; for each action there is a ‘common’ (high probability) transition and an ‘uncommon’ (low probability) one. Each second-stage state then delivers a reward with a particular probability, which changes intermittently (see Methods).

**Figure 5.**
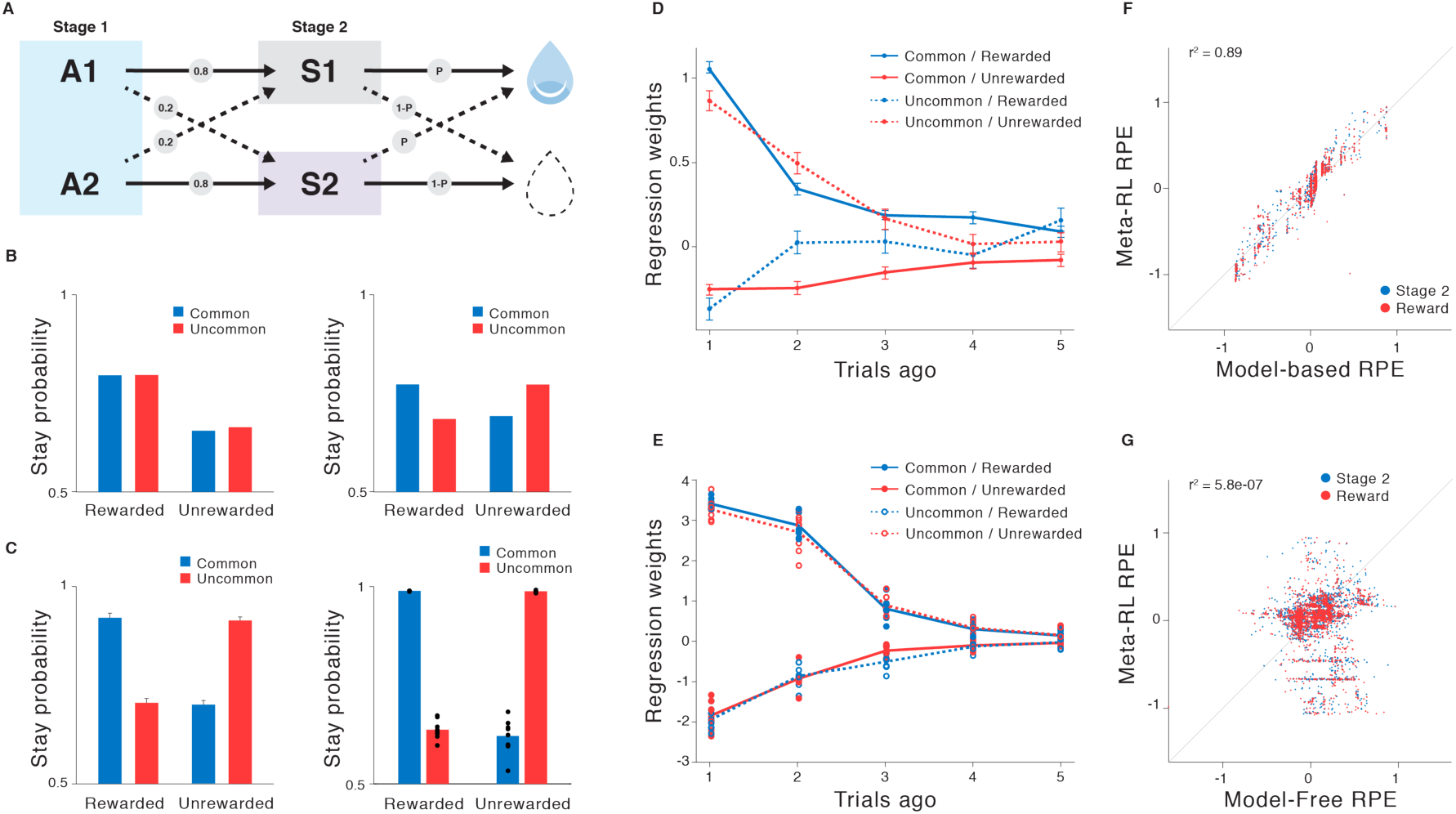
A. Structure of the two-step task.^21,55^ B. Top: Canonical pattern of behavior for model-free (left) and model-based (right) learning. C. Left: Data from Miller et al.^55^ showing a canonical model-based pattern in animal behavior. Plot represents mean stay probability for n = 6 rats, error bars are s.e.m. Right: Corresponding behavior of the network. Dots represent data from separate training runs using different random seeds (n = 8). D. More detailed analysis from Miller et al., showing the influence of transition types and rewards, at multiple trial lags, on animal behavior. The pattern rules out an alternative, model-free mechanism (see Supplementary Figure 4). E. Corresponding analysis of meta-RL behavior. Dots represent data from separate training runs using different random seeds (n = 8). F, G. Regression of model RPE signals against predictions from model-based (F) and model-free (G) algorithms. Analysis done for 30 evaluation episodes, each consisting of 100 trials, performed by 1 fully trained network out of 8 replicas. Panels C (left) and D adapted with permission from Miller et al.^55^

The interest of the two-step task lies in its ability to differentiate between model-free and model-based learning. Most important is behavior following an uncommon transition. Consider an action selected at stage one which triggers an uncommon transition followed by a reward. Model-free learning, driven by action-reward associations, will increase the probability of repeating the same first-stage action on the subsequent trial. Model-based learning, in contrast, will take account of the task’s transition structure and increase the probability of choosing the opposite action (Figure 5B). Both humans^21^ and rodents^55^ typically display behavioral evidence of model-based learning (see Figure 5C,D).

We trained our meta-RL model on the two-step task (see Methods), and found that its behavior at test assumed the form associated with model-based control (Figure 5C,E). As in previous simulations, the network was tested with weights frozen. The observed behavior was thus generated by the dynamics of the recurrent prefrontal network. However, it is important to recall that the network was trained by a *model-free* DA-driven RL algorithm. In a neuroscientific context, this points to the interesting possibility that the PFC’s implementation of model-based RL might in fact arise from model-free DA-driven training (see Supplementary Figures 4–6 for further analysis).

Critically, the two-step task was one of the first contexts in which structure-sensitive RPE signals were reported. In particular, using fMRI, Daw and colleagues^21^ observed RPEs in human ventral striatum — a major target for DA — which tracked those predicted by a model-based RL algorithm. The same effect arises in our meta-RL model, as shown in Figure 5F,G.

### Simulation 5. Learning to learn

Simulations 3 and 4 focused on scenarios involving an alternation between two versions of a task, each of which might become familiar over time. Here we apply meta-RL to a task in which new stimuli are continually presented, requiring learning in the fullest sense. In this setting, we show that meta-RL can account for situations where past experience speeds new learning, an effect often called “learning to learn.”

In the task that originally inspired this term, Harlow^56^ presented a monkey with two unfamiliar objects, one covering a well containing food reward, the other an empty well. The animal chose freely between the objects and could retrieve the food reward if present. The left-right positions of the objects were then randomly reset, and a new trial began. After six repetitions of this process, two entirely new objects were substituted, and the process began again. Importantly, within each block of trials, one object was chosen to be consistently rewarded, with the other consistently unrewarded. Early in training, monkeys were slow to converge on the correct object in each block. But after substantial practice, monkeys showed perfect performance after only a single trial, reflecting an understanding of the task’s rules (Figure 6B).

**Figure 6.**
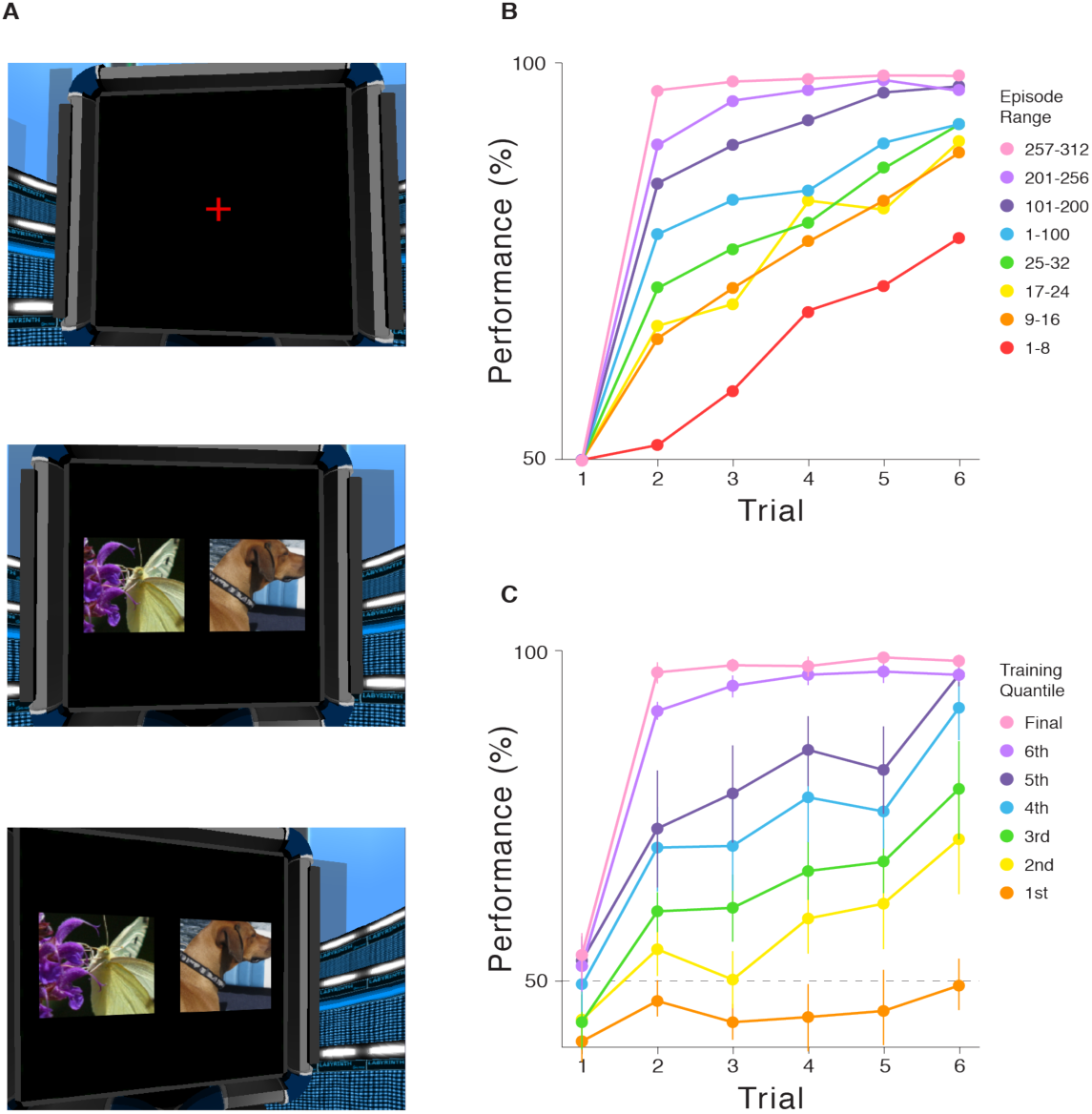
A. Example image inputs for Simulation 5 (full resolution), showing fixation cross (top), initial stimulus presentation (middle) and saccade outcome (bottom). B. Accuracy data from Harlow^56^ for each step following introduction of new objects and at multiple points in training of connection weights. C. Model performance at seven successive stages of training (see Methods), averaged across all networks (n = 14) that achieved maximal performance (out of 50 replicas). Error bars represent 95% CIs.

To evaluate the ability of meta-RL to account for Harlow’s results, we converted his task to one requiring choices between images presented on a simulated computer display (Figure 6A; Supplementary Video 1; https://youtu.be/FXxAobxqPUo). The task was otherwise unchanged, with a pair of unfamiliar images introduced after every six trials. To allow our model to process high-dimensional pixel inputs, we augmented it with a conventional image-processing network (see Methods). The resulting system generated learning curves closely resembling the empirical data (Figure 6C). Following training, the network learned in ‘one shot’ how to respond in each new block, replicating Harlow’s learning-to-learn effect.

### Simulation 6. The role of dopamine: Effects of optogenetic manipulation

The meta-RL framework requires that the inputs to the prefrontal network contain information about recent rewards. Our implementation satisfies this requirement by feeding in a scalar signal explicitly representing the amount of reward received on the previous time-step (see Figure 1A). However, any signal that is robustly correlated with reward would suffice. Sensory signals, such as the taste of juice, could thus play the requisite role. Another particularly interesting possibility is that information about rewards might be conveyed by DA itself. In this case, DA would play two distinct roles. First, as in the standard model, DA would modulate synaptic plasticity in the prefrontal network. Second, the same DA signal would support activity-based RL computations within the prefrontal network, by injecting information about recent rewards^57^. We modified our original network to provide the RPE, in place of reward, as input to the network (see Methods and Supplementary Figure 7), and found that it generated comparable behavior on the tasks from Simulations 1–5.

This variation on meta-RL suggests a novel interpretation for the findings of recent experiments using optogenetic techniques to block or induce dopaminergic RPE signals^58–61^. To illustrate, we consider an experiment by Stopper and colleagues^62^, involving a two-armed bandit task with intermittently shifting payoff probabilities (see Methods). Blocking DA activity during delivery of food rewards from one lever led to a reduced preference for that lever. Conversely, artificially stimulating DA when one lever failed to yield food increased preference for that lever (Figure 7A). We simulated these conditions by incrementing or decrementing the RPE input to the network in Supplementary Figure 7, and observed comparable behavioral results (Figure 7B). As in previous simulations, these results were obtained with the network weights held constant. The shift in behavior induced by our simulated optogenetic intervention thus reflects the impact of the dopaminergic RPE signal on unit activities within the prefrontal network, rather than an effect on synaptic weights. In this regard, the present simulation thus provides an interpretation of the experimental results that is radically different from the one that would proceed from the standard model.

**Figure 7.**
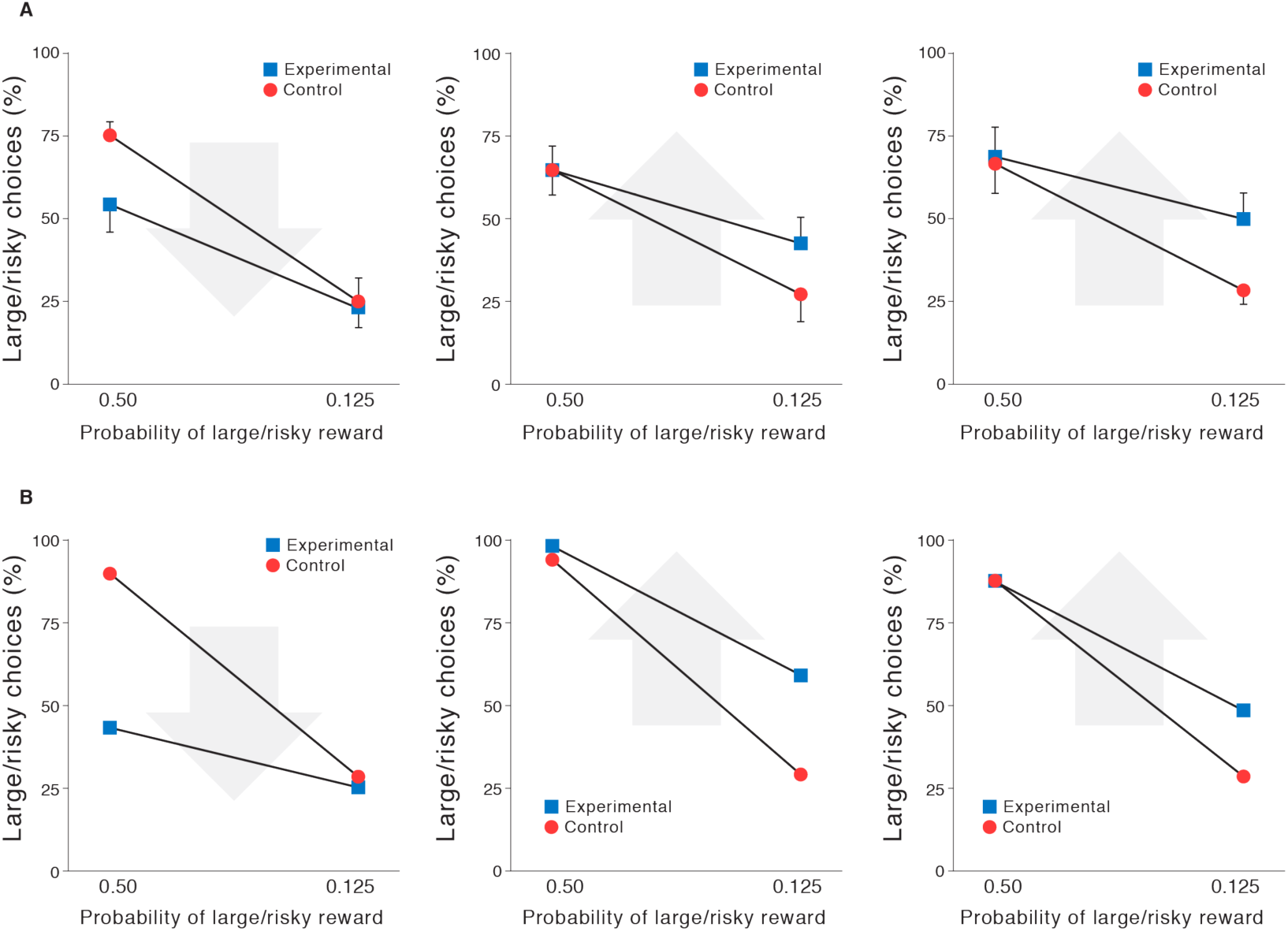
A. Behavioral data adapted with permission from Stopper et al.^62^. Their task posed a recurring choice between a lever that reliably yielded a small reward (small/certain) and one that yielded a large reward with probability *p*(large/risky). In a control condition, rats chose the large/risky lever more often when *p* = 0.5 than when *p* = 0.125 (blue). Left: Effect of optogenetically blocking DA when large/risky rewards occurred (n = 11). Arrow indicates direction of shift with optogenetic intervention. Center: Effect of blocking DA when small/certain rewards occurred (n = 8). Right: Effect of triggering DA when large/risky rewards failed to occur (n = 9). All plots represent mean ± s.e.m. B. Model behavior in circumstances modeling those imposed by Stopper et al.^62^, with conditions presented as in A.

### Functional Neuroanatomy

As our simulation results show, meta-RL provides a new lens through which to examine the respective functions the prefrontal network and the DA system, and their interactions during learning.

As we have noted, a key assumption of the meta-RL theory is that the prefrontal network can be understood as a recurrent neural circuit^23–26^. In fact, the PFC lies at the intersection of multiple recurrent circuits, involving recurrent connections within PFC itself, connections with other cortical regions^63^ and, most crucially, connections with dorsal striatum and mediodorsal thalamus (see Supplementary Figure 8). A prevalent view^24^ is that the role of the striatum within this cortico-basal ganglia-thalamo-cortical loop is to regulate the dynamics of PFC by gating the flow of information into PFC circuits. According to this account, DA serves as a training signal within striatum, shaping the operation of the gating function through TD learning. Replicating our simulation results using this more detailed model of the cortico-striatal loop represents a worthwhile target for next-step research.

It is worth noting that the striatal gating theory was originally inspired by long short-term memory (LSTM) networks, of the kind we have employed in our simulations^24^ (see Figure 1B). Indeed, recent work posits multiple gating mechanisms operating on PFC, which directly parallel the input, memory and output gating mechanisms traditionally implemented in LSTM networks^64^ (see Methods). While these parallels are intriguing, it should also be noted that other recent work has modeled RL within the PFC using more generic recurrent neural networks that lack the gating mechanisms involved in LSTM networks^23, 25^. Although the meta-reinforcement learning effect we have identified relies only on the presence of recurrence, and not on any particular gating mechanism, it would be very interesting to test whether particular gating mechanisms provide a more precise quantitative fit to human and animal learning curves and neural representations.

The neuroscientific data we have considered in this paper center mostly on structures within the so-called ‘associative loop’ connecting the dorsal PFC with the basal ganglia^65^. However, this is only one of several such recurrent loops in the brain, with others running through other sectors of cortex^65^ (see Supplementary Figure 8). This raises the question of whether other recurrent loops, such as the ‘sensorimotor loop’ running through somatosensory and motor cortices, also support meta-RL. Although we cannot rule out that forms of meta-learning may also emerge in these loops, we observe that the meta-RL effect described in this paper emerges only when network inputs carry information about recent actions and rewards, and when network dynamics support maintenance of information over suitable time-periods. These two factors may differentiate the associative loop from other cortico-basal ganglia-thalamo-cortical loops^66^ explaining why meta-RL might arise uniquely from this circuit (see the Supplementary Figure 8).

Most of our simulations modelled all of PFC as a single fully-connected network without regional specialization. However, there are important functional-anatomical distinctions within PFC. For example, the probability-matching behavior seen in Simulation 1 and the model-based learning pattern in Simulation 4 have robust neural correlates in DLPFC^10, 17^, while the volatility coding modelled in our Simulation 2 was reported in the anterior cingulate cortex^46^. Although the differential roles of anterior cingulate and DLPFC are still under active debate, an important next step for the present theory will be to consider how the computations involved in meta-RL might play out across these regions, given their different cell properties, internal circuitry and extrinsic connectivity. Also relevant are PFC regions lying within the so-called ‘limbic loop’^65^, including orbitofrontal and ventromedial PFC (see Supplementary Figure 8). Both of the latter regions have been implicated in reward coding, while recent work suggests that the orbitofrontal cortex may additionally encode abstract latent states^49, 50^, both critical functions for meta-RL, as illustrated in several of our simulations. Once again, a fuller development of the meta-RL theory will need to incorporate the relative roles of these regions more explicitly.

## Discussion

We have put forward a new proposal concerning the roles of DA and PFC in reward-based learning, leveraging the notion of meta-reinforcement learning. The framework we have advanced conserves the standard RPE model of DA function but places it in a new context, allowing it to newly accommodate a number of previously puzzling findings. As our simulations have shown, meta-RL accounts for a diverse range of observations concerning both DA and PFC function, providing a bridge between the literatures addressing these two systems.

In addition to explaining existing data, the meta-RL framework also leads to a number of testable predictions. As we have seen, the theory suggests that the role of PFC in model-based control could arise, at least in part, from DA-driven synaptic learning. If this is correct, then interfering with phasic DA signaling during initial training should interfere with the emergence of model-based control in tasks like those studied in Simulations 4–5. Another prediction concerns model-based dopaminergic RPE signals. Meta-RL attributes these signals to value inputs from the prefrontal network. If this is correct, then lesioning or inactivating PFC or its associated striatal nuclei should eliminate model-based DA signaling^67^. Further predictions might be specified by examining the patterns of activity arising in the prefrontal network portion of our model, treating these as predictors for neural activity in animals performing relevant tasks. Behavioral paradigms useful to such an undertaking might be drawn from recent work implicating PFC in the identification of latent states^27, 49–54^ and abstract rules^68, 69^ in RL contexts.

Meta-RL also raises a range of broader questions which we hope will stimulate new experimental work. What might be the relative roles of mesolimbic, mesocortical and nigrostriatal DA pathways, in a meta-RL context? Does meta-RL point to new interpretations of the division of labor between dorsal and ventral, or medial and lateral sectors of PFC? How should we interpret data pointing to the existence of systems supporting both model-based and model-free RL (see Supplementary Figures 5, 6)? What new dynamics emerge when meta-RL is placed in contact with mechanisms subserving episodic memory^70, 71^? All in all, meta-RL offers a new orienting point within the landscape of ideas concerning reward-based learning, one that may prove useful in the development of new research questions and in the interpretation of new findings.

## Methods

### Architecture and Learning Algorithm

All of our simulations employed a common set of methods, with minor implementational variations. The agent architecture centers on a fully connected, gated recurrent neural network (LSTM: long short-term memory network^72^, see below for equations). In all experiments except where specified, the input included the observation, a scalar indicating the reward received on the preceding time-step, and a one-hot representation of the action taken on the preceding time-step. The outputs consisted of a scalar baseline (value function) and a real vector with length equal to the number of available actions. Actions were sampled from the softmax distribution defined by this vector. Some other architectural details were varied as required by the structure of different tasks (see simulation-specific details below). Reinforcement learning was implemented by Advantage Actor-Critic, as detailed in Mnih et al.^43^. Details of training, including the use of entropy regularization and a combined policy and value estimate loss, are described in Mnih et al.^43^ In brief, the gradient of the full objective function is the weighted sum of the policy gradient, the gradient with respect to the state-value function loss, and an entropy regularization term, defined as follows:

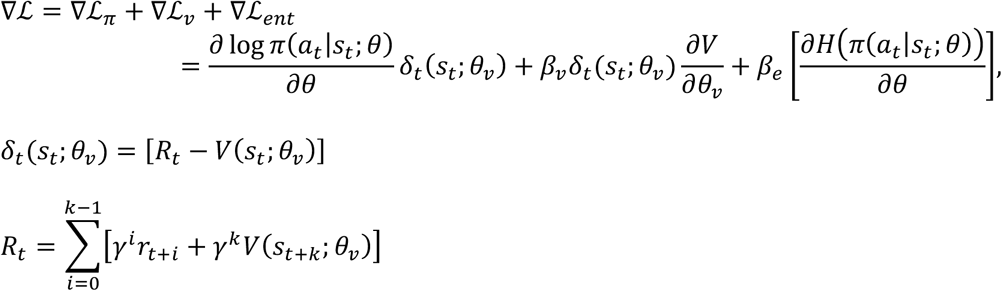

where *a_t_, s_t_*, and *R_t_* define the action, state, and discounted *n*-step bootstrapped return at time *t* (with discount factor *γ*), *k* is the number of steps until the next terminal state and is upper bounded by the maximum unroll length *t_max_, π* is the policy (parameterized by neural network parameters *θ*), *V* is the value function (parameterized by *θ_v_*), estimating the expected return from state *s, H*(*π*) is the entropy of the policy, and *β_v_* and *β_e_* are hyperparameters controlling the relative contributions of the value estimate loss and entropy regularization term, respectively. *δ_t_*(*s_t_*; *θ_v_*) is the *n*-step return TD-error that provides an estimate of the advantage function for actor-critic. The parameters of the neural network were updated via gradient descent and backpropagation through time, using advantage actor-critic as detailed in Mnih et al.^43^ Note that while the parameters *θ* and *θ_v_* are being shown as separate, as in Mnih et al.^43^, in practice they share all non-output layers and differ only in the softmax output for the policy and one linear output for the value function. Simulations 1–4 and 6 used a single thread and received discrete observations coded as one-hot vectors, with length of the number of possible states (see Algorithm 1 for pseudocode for single-threaded advantage actor-critic). Simulation 5 used 32 asynchronous threads during training and received RGB frames as input (see Mnih et al.^43^ for asynchronous multi-threaded algorithm and pseudocode). The core recurrent network consisted of 48 LSTM units in simulations 1–4, 256 units in simulation 5, and two separate LSTMs of 48 units each (for policy and value) in simulation 6 (see below).

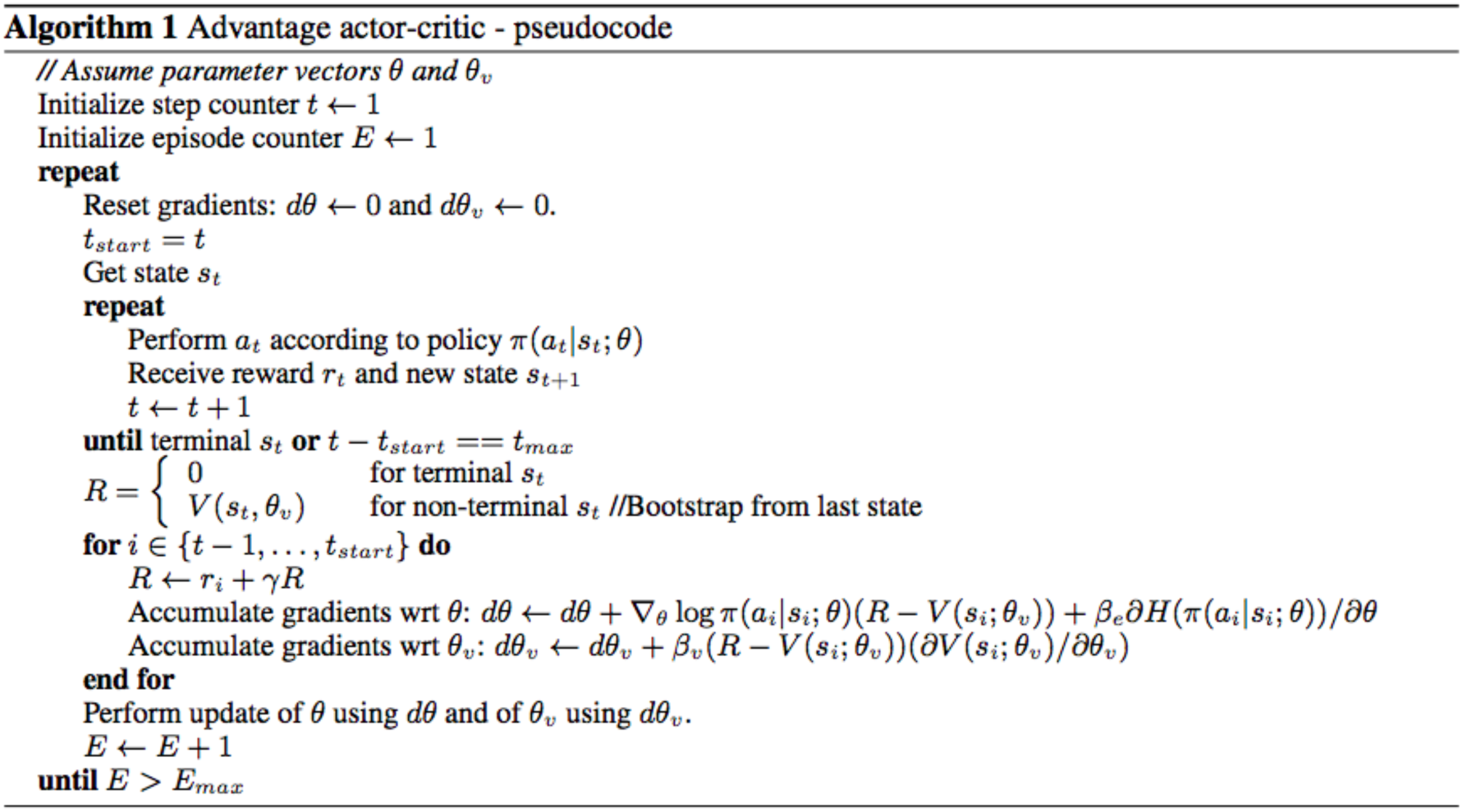

In a standard non-gated recurrent neural network, the state at time step *t* is a linear projection of the state at time step *t*–1, followed by a nonlinearity. This kind of “vanilla” RNN can have difficulty with long-range temporal dependencies because it has to learn a very precise mapping just to copy information unchanged from one time state to the next. An LSTM, on the other hand, works by copying its internal state (called the “cell state”) from each time step to the next. Rather than having to learn how to remember, it remembers by default. However, it is also able to choose to forget, using a “forget” (or maintenance) gate, and to choose to allow new information to enter, using an “input gate”. Because it may not want to output its entire memory contents at each time step, there is also an “output gate” to control what to output. Each of these gates are modulated by a learned function of the state of the network.

More precisely, the dynamics of the LSTM were governed by standard equations^72, 73^:

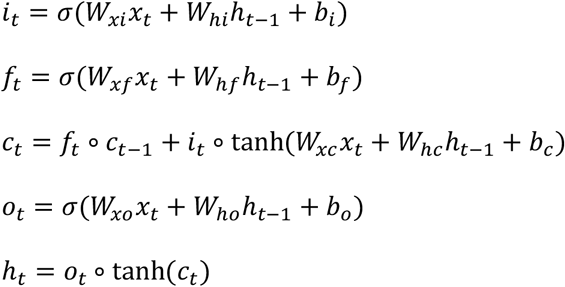

where *x_t_* is the input to the LSTM at time *t, h_t_* is the hidden state, *i_t_* is the input gate, *f_t_* is the forget/maintenance gate, *o_t_* is the output gate, *c_t_* is the cell state, *σ* is the sigmoid function, and ∘ is an operator denoting element-wise multiplication.

### General task structure

Tasks were episodic, comprising a set number of trials (fixed to a constant number per episode unless otherwise specified), with task parameters randomly drawn from a distribution and fixed for the duration of the episode. In simulations 1, 3, 4 and 6, each trial began with a fixation cue, requiring a distinct central fixation response (*a_c_*), followed by one or more stimulus cues, each requiring a stimulus response (either left, *a_l_*, or right, *a_r_*). Failure to produce a valid response to either the fixation or stimulus cues resulted in a reward of −1. In simulation 5, which involved high dimensional visual inputs, fixation and image selection required emitting left-right actions continuously to shift the target to within the center of the field of view. An additional no-op action was provided to allow the agent to maintain fixation as necessary, and no negative reward was given for producing invalid actions. See simulation-specific methods for more details.

### Training and testing

Both training and testing environments involved sampling a task from predetermined task distributions — in most cases sampling randomly, although see Simulations 1 and 2 for principled exceptions — with the LSTM hidden state initialized at the beginning of each episode (initial state learned for simulations 1–4 and 6; initialized to 0 for simulation 5). Unless otherwise noted, training hyperparameters (as defined in Mnih et al.^43^) were as follows: learning rate = 0.0007, discount factor = 0.9, state-value estimate cost *β_v_* = 0.05, and entropy cost *β_e_* = 0.05. Weights were optimized using Shared RMSProp and backpropagation through time^43^, which involved unrolling the recurrent network a fixed number of timesteps that ranged from *t_max_* = 100–300 depending on task, and determined the number of steps when calculating the bootstrapped *n*-step return. The agent was then evaluated on a testing episodes, during which all network weights were held fixed. No parameter optimization was undertaken to improve fits to data, beyond the selection of what appeared to be sensible *a priori* values, based on prior experience with related work, and minor adjustment simply to obtain robust task acquisition.

We now detail the simulation-specific task designs, hyperparameters, and analyses. Note that details of the simulations reported in the introductory two-armed bandit simulations were drawn from Wang et al.^41^.

#### Simulation 1. Reinforcement learning in the prefrontal network

This task, from Tsutsui et al.^10^, required the agent to select between two actions, *a_l_* and *a_r_*, with dynamic reward probabilities that changed in response to previous choices. Reward probabilities for actions were given according to

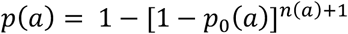

The value of *p_0_(a)*, the baseline probability of reward for action *a*, was sampled from a uniform Bernoulli distribution and held fixed for the entire episode. *n(a)* was the number of trials since action *a* was chosen. The sum of the baseline probabilities for both arms was always constant and equal to 0.5, so that *p_0_ (a_L_)* = 0.5 – *p_0_ (a_R_)*. The number of trials per episode varied uniformly from 50 to 100 (chosen randomly at the beginning of the episode). For training only, we held out the subset of environmental parameters *p*_0_ = 0.1 - 0.2 and 0.3 - 0.4, so the agent was only trained on *p_0_* = [0.0-0.1]∪[0.2-0.3]∪[0.4-0.5]. For this task, we used discount factor = 0.75 and trained for 3e6 steps (∼15,000 episodes).

After training, we tested the agent for 500 episodes with network weights held fixed. Analyses of model behavior were based on the original study, which in turn employed techniques introduced by Lau and Glimcher^44^. We determined the influence of past choices and reward outcomes on trial-by-trial responses by fitting to a logistic regression model, as in Lau and Glimcher^44^ and Tsutsui et al.^10^, using a maximum trial lag of *N* = 15. Only trials from the last two-thirds of each episode were considered, in order to restrict our analyses to steady-state behavior.

To assess the amount of information encoded about each variable in the recurrent network, we calculated Spearman’s partial rank correlation between hidden unit activations and four factors: the last action, last reward, action × reward interaction, and choice value. (Note that our task implementation abstracted over the distinction between actions and target objects, and we do not focus on the distinction between action and object value.) Because the values of the two actions were strongly anti-correlated (*r* = −0.95, Pearson’s correlation), we only considered the effect of the left action.

#### Simulation 2. Adaptation of prefrontal-based learning to the task environment

On each step the agent chose between two actions, *a_l_* and *a_r_*, and received a reward of *R* or 0. The magnitudes of *R* that could be obtained varied randomly across trials and independently between the two actions (distributed as 100*Beta(4,4)), and were cued to the agent on each trial. The reward probabilities of the two arms were perfectly anticorrelated and thus described by a single parameter r(t), which evolved over time. In testing, the evolution of r(t) mirrored the task human subjects performed in Behrens et al.^46^. There were two types of episodes. In episodes with low volatility first, r(t) remained stable at 0.25 or 0.75 for 100 trials, and then alternated between 0.2 and 0.8 every 25 trials. In episodes with high volatility first, r(t) alternated between 0.2 and 0.8 every 25 trials for 100 trials, and then remained stable at 0.25 or 0.75.

For training, we adapted the two-level generative model used for inference in Behrens et al. Specifically, in training, r(t) evolved according to v(t), such that with probability v(t), r(t+1) was set to 1-r(t), and with probability 1-v(t), r(t+1) remained the same as r(t). Analogously, the volatility v(t) also evolved over time. With probability k, v(t+1) was set to 0.2-v(t), and with probability 1-k, v(t+1) remained the same as v(t), where log(k) was sampled uniformly at the end of each episode between −4.5 and −3.5. At the beginning of each training episode, v(1) was randomly initialized to 0 or 0.2. When v(t) was 0, r(t) was set to 0.25 or 0.75, and when v(t) was 0.2, r(t) was set to 0 or 1. Tying r(t) to v(t) in this way matched the structure of the task subjects performed in Behrens et al.^46^ (see above and Figure 3A). The probabilistic model employed in that study did not impose this connection, with the result that dynamic adjustments in learning rate had less effect on expected reward. For completeness, we also ran simulations in which meta-RL was trained using data produced from the generative model employed by Behrens et al.^46^, and obtained qualitatively similar results (data not shown).

Meta-RL was trained for 40,000 episodes and tested for 400 episodes, consisting of 200 trials each. Because of the length of these episodes, we did not include a fixation step. Training was conducted with learning rate = 5e-5, discount = 0.1, and an entropy cost linearly decreasing from 1 to 0 over the course of training. As in all simulations, weights were held fixed in testing.

For purposes of analysis, following Behrens et al.^46^, we implemented a Bayesian agent, which performed optimal probabilistic inference under the generative model that defined the training experience. The Bayesian agent used a discretized representation of the posterior as in Behrens et al.^46^. The “estimated volatility” signal we report was the agent’s expectation of volatility at time *t*. We also fit a multiple regression model to predict the Bayesian agent’s estimated volatility at each time step from the recurrent network’s hidden activations, and applied this model to hidden activations on held-out episodes (in cross-validation) to generate the “decoded volatility” signal we report for meta-RL.

Learning rates were also estimated for both LSTM and Bayes, based only on their behavior. To estimate learning rates, we fit a Rescorla-Wagner model to the behavior by maximum likelihood. Episodes of the two types (high volatility first and low volatility first) were fit separately. The Rescorla-Wagner model was fit to behavior from all episodes, but from only 10 consecutive trials, sliding the window of 10 trials in single-trial increments to observe how learning rates might change dynamically. Although only 10 consecutive trials contributed directly to the likelihood, all trials previous to these 10 nevertheless contributed indirectly through their influence on the evolving value estimate. As a control, we estimated learning rates for behavior generated by a Rescorla-Wagner model with a fixed learning rate, to ensure we could recover the true learning rate without bias.

To determine the proportion of hidden units coding for volatility, in each episode we regressed the activity of each hidden unit against the true volatility, coded as 0 or 1. (This regression model also included several covariates: previous reward, next action, and true probability of reward on the present time step. However, including these covariates had little effect on the proportion of units discovered as coding volatility.) We then counted units for which the slope of the volatility regressor was significantly different from zero, at a threshold of *p* = 0.05, Bonferroni corrected for number of hidden units. The mean and standard error of this number are reported in the main text.

In a similar analysis, we also regressed across episodes at each time step. In Supplementary Figure 3A we plot over time the proportion of units for which the slope of the volatility regressor was significantly different from zero, at a Bonferroni corrected threshold of *p* = 0.05. In Supplementary Figure 3B we continuously vary the threshold across a wide range and plot the proportion of units, averaged over trials, exceeding the threshold.

Finally, we note that in Behrens et al (2007) the estimated volatility signal from the Bayesian model was used as a regressor, while we use the true volatility. We found that these two approaches did not produce qualitatively different results. However, it was slightly more difficult to interpret the Bayes estimated volatility signal at a fine-grained scale of individual trials, because the Bayes estimated signal was sensitive to individual trial events. Particularly early within an episode, the Bayes estimated signal was more a reflection of the specific outcomes that had been received than a reflection of volatility.

#### Simulation 3. Reward prediction errors reflecting inferred value

Following Bromberg-Martin et al.^18^, we trained our agent on a reversal task, in which the values of the two stimulus cues were anti-correlated, with one being rewarded and the other not rewarded. The rewarded stimulus switched intermittently in an uncued manner, such that there was a 50% chance of a switch at the beginning of each episode. Concretely, after the central fixation cue on step 1, the agent was presented with one of the two stimulus cues (tabular one-hot vectors) on step 2, indicating that it must either produce an action left (*a_l_*) or an action right (*a_r_*) on step 3. The reward was then delivered on step 4, followed by the start of the next trial. Failure to produce a valid response on any step gave a reward of −1, while a valid action resulted in a reward of 1 for the rewarded cue, or 0 for the unrewarded cue and central fixation cue.

The rewarded and unrewarded cues were randomly determined at the beginning of the episode and held fixed for the duration of the episode, lasting 5 trials (20 steps). After training for 2e6 steps, or 1e5 episodes, using a backpropagation window of 200 steps, we tested for 1200 episodes and calculated the reward prediction errors in response to stimulus presentation (step 2) for trials 1 and 2:

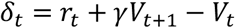

Baselines (i.e. respective *V_t_*) were also extracted from steps 1 and 3 for comparison. We analyzed all test episodes in which a reward reversal had occurred with respect to the previous episode.

#### Simulation 4. ‘Model-based’ behavior: The Two-Step Task

The structure of the two-step task, diagrammed in Figure 5A, was based directly on the version used by Miller et al.^55^. After central fixation, on the first stage of the trial the agent chose either *a_l_* or *a_r_* and transitioned into one of two second-stage states *S_1_* or *S_2_* with probabilities *P(S_1_/a_L_)* = *P(S_2_/a_R_)* = 0.8 (*common* transition) and *P(S_1_/a_R_)* = *P(S_2_/a_L_)* = 0.2 (*uncommon* transition). The transition probabilities *P* were fixed across episodes. The states *S_1_* and *S_2_* yielded probabilistic reward according to [*P(r/S_1_), P(r/S_2_)*] = [0.9, 0.1] or [0.1, 0.9], with the specific reward contingencies having a 2.5% chance of randomly switching at the beginning of each trial. The agent was trained for 10,000 episodes of 100 trials each, and tested with weights fixed on 300 further episodes. Behavioral analyses (results shown in Figure 5C,E) closely followed corresponding procedures specified in Miller et al.^55^ and Daw et al.^21^.

To compare the RPEs of our meta-RL agent to RPEs produced by model-free and model-based algorithms, we followed the procedure employed by Daw et al.^21^. As in that study, the reference model-free algorithm was SARSA(0) and the model-based algorithm was a prospective planner, both as implemented in Daw et al.^21^. However, instead of performing Q-learning at the second stage, which is not optimal, the model-based algorithm tracked the latent variable H describing whether *P(r|S_1_)* was 0.9 (equivalent to whether *P(r|S_2_)* was 0.1). The algorithm updated its belief about H following each outcome, using knowledge of the task’s true structure. Both algorithms were applied to the same trial sequence experienced by the meta-RL agent. Each agent generated a reward prediction error on each step, as described in Daw et al.^21^, but with the expectation of reward at the second stage being *P(H)* * 0.9 + (1 – *P*(*H*))* 0.1 if the agent was in state S_1_ and (1 - *P*(*H*)) * 0.9 + *P*(*H*) * 0.1 if the agent was in state S_2_.

RPEs from meta-RL were calculated as in Simulation 3. We were interested in determining whether meta-RL’s RPEs were model-based only, model-free only, or a mixture of both. This analysis must take account of the fact that model-free and model-based RPEs themselves are somewhat correlated (Supplementary Figure 4). We first regressed meta-RL’s RPEs directly against the model-based RPEs of the prospective planner. Finding an almost perfect correlation, we then asked whether meta-RL’s RPEs were at all related to the component of model-free RPEs that is orthogonal to the model-based RPEs. We did this by regressing the model-based RPE out of the model-free RPE and looking for a correlation of meta-RL’s RPE with the residual. We note that, unlike subjects in the fMRI study by Daw and colleagues, our model showed a purely model-based pattern of behavior (see Figure 5C,E). Correspondingly, we found exclusively model-based RPEs in the present analysis. As a confirmatory analysis, we also performed a multiple regression with meta-RL’s RPEs as the dependent variable, and both model-based and model-free RPEs as independent variables (Supplementary Figure 4).

#### Simulation 5. Learning to learn

An earlier version of this simulation was presented by Wang et al.^41^. To simulate Harlow’s^56^ learning-to-learn task, we trained our agent with RGB pixel input using the PsychLab framework described by Leibo et al. (2018).^74^ An 84×84 pixel input represented a simulated computer screen (see Figure 6A and Supplementary Video 1; https://youtu.be/FXxAobxqPUo). At the beginning of each trial, this display was blank except for a small central fixation cross. The agent selected discrete left-right actions which shifted its view approximately 4.4 degrees in the corresponding direction, with a small momentum effect (alternatively, a no-op action could be selected). The completion of a trial required performing two tasks: saccading to the central fixation cross, followed by saccading to the correct image. If the agent held the fixation cross in the center of the field of view (within a tolerance of 3.5 degrees visual angle) for a minimum of four time steps, it received a reward of 0.2. The fixation cross then disappeared and two images — drawn randomly from the ImageNet dataset^75^ and resized to 34×34 — appeared on the left and right side of the display (see Figure 6A). The agent’s task was then to “select” one of the images by rotating until the center of the image aligned with the center of the visual field of view (within a tolerance of 7 degrees visual angle). Once one of the images was selected, both images disappeared and, after an intertrial interval of 10 time-steps, the fixation cross reappeared, initiating the next trial. Each episode contained a maximum of 6 trials or 3600 steps. Each selected action was mandatorily repeated for a total of four time-steps (as in Mnih et al.^43^), meaning that selecting an image took a minimum of three independent decisions (twelve primitive actions) after having completed the fixation. It should be noted, however, that the rotational position of the agent was not limited; that is, 360 degree rotations could occur, while the simulated computer screen only subtended 65 degrees.

Although new ImageNet images were chosen at the beginning of each episode (sampled with replacement from a set of 1000 images), the same two images were re-used across all trials within an episode, though in randomly varying left-right placement, similar to the objects in Harlow’s experiment. And as in that experiment, one image was arbitrarily chosen to be the “rewarded” image throughout the episode. Selection of this image yielded a reward of 1.0, while the other image yielded a reward of −1.0. The LSTM was trained using backpropagation through time with 100-step unrolls.

During test, network weights were held fixed (i.e. in convolutional network and LSTM-A3C) and ImageNet images were drawn from a separate held-out set of 1000, never presented during training. Learning curves were calculated for each replica by taking a rolling average of returns over episodes and normalizing from 0 to 100%, where 50% represented performance of randomly choosing between the two images and 100% represented optimal performance. We trained 50 networks using hyperparameters adopted from Wang et al.^41^ (learning rate = 0.00075, discount = 0.91, entropy cost *β_e_* = 0.001, and state-value estimate cost *β_v_* = 0.4) and found that 28% reached maximal performance after ∼1e5 episodes (per thread, 32 threads). Because this task required not only learning abstract rule structure, but also processing visually complex images, staying oriented toward the simulated computer screen, and central fixation, much of the initial training time was spent at 0% performance. Ultimately successful agents varied in the number of episodes required to overcome these initial hurdles (median 3413, range 1810–29,080). To characterize acquisition of abstract task structure dissociated from these other considerations, we identified two change-points in the learning curves (achieving above-chance level performance and approaching ceiling performance) and plotted average reward as a function of trial number within the episode at various quantiles of performance interpolated between these change-points (6 quantiles) as well as the final performance at end of training. Agents also varied in the number of episodes required to achieve maximal performance after surpassing chance-level performance (median 4875, range 1500–82,760).

In order to be sure that the meta-RL network was truly performing the role-filler assignment that is intended to be required by the Harlow task, rather than relying on a simpler strategy, we trained the same model on a variant of the Harlow task where the non-target (unrewarded) image changed across trials, always taking the form of a novel, unfamiliar picture. Following training, the performance of our network was identical to its performance in the original task, with chance accuracy on trial one followed by near ceiling accuracy for the remainder of the same trial block (data not shown).

#### Simulation 6. The role of dopamine: Effects of optogenetic manipulation

We implemented a probabilistic risk/reward task as described in Stopper et al.^62^, in which subjects chose between a “safe” arm that always offered a small reward (*r_s_* = 1) or a “risky” arm that offered a large reward (*r_l_* = 4) with a probability *P*(*a_Large_*) = 0.125 (Safe Arm Better block) or 0.5 (Risky Arm Better block), sampled at the beginning of the episode. In direct analogy with the original study, the agent was first required to make 5 forced pulls each of the safe and risky arms (in randomized pairs), followed by 20 free pulls. The locations (i.e. associated actions) of the risky and safe arms could switch from episode to episode, with a probability of 0.5.

In order to simulate optogenetic stimulation, it was necessary to implement a variant of our original architecture in which two separate LSTMs (48 units each) model value estimate and policy (rather than a single LSTM with two linear outputs), in alignment with standard actor/critic architectures associated with basal ganglia and PFC function (see Supplementary Figure 8). Concretely, the critic is modeled as an LSTM which takes as input the state observation, last reward received, and last action taken, outputting value estimate. The actor receives the state observation, last action taken, and the reward prediction error that is computed based on the value estimate output from the critic, and subsequently outputs the policy (see Supplementary Figure 7). Because the RPE is computed based on the prefrontal network value output, it could not be fed as an input to the critic, given our other architectural and algorithmic assumptions.

Optogenetic stimulation was simulated by manipulating the value of the reward prediction error fed into the actor, in directly analogy with lateral habenula and VTA stimulation in Stopper et al.^62^. After training our agent in the control condition in which normal RPEs were input to the actor (4e6 steps; 6.67e4 episodes of 30 trials each), we tested our agent for 1333 episodes in four different conditions designed to approximate the different stimulation protocols employed in Stopper et al., 2014: 1) control (analogous to Stopper’s baseline no stimulation condition), 2) block risky reward - if the risky arm was chosen and rewarded, subtract 4 from the RPE (approximating LHb stimulation with 4 trains of pulses), 3) block safe reward - if the safe arm was chosen, subtract 1 from the RPE (approximating LHb stimulation with 1 train of pulses), and 4) block risky loss - if the risky arm was chosen and not rewarded, add a reward of 1 to the RPE (approximating VTA stimulation).

## Acknowledgements

We are grateful to K. Miller, F. Grabenhorst, T. Behrens, E. Bromberg-Martin, S. Floresco, and P. Glimcher for graciously providing help with and permission for adapting their data. We thank C. Blundell and R. Munos for useful discussions and comments on an earlier draft of the paper. The opinions expressed in this publication are those of the authors and do not necessarily reflect the views of the funding agencies.

## Author Contributions

J.X.W., Z.K.N., and M.B. designed the simulations. J.W. and Z.K.N. performed the simulations and analyzed the data. D.T., H. S., J.Z.L. contributed and helped with code. All authors wrote the manuscript.

## Supplementary Figures

**Supplementary Figure 1.**
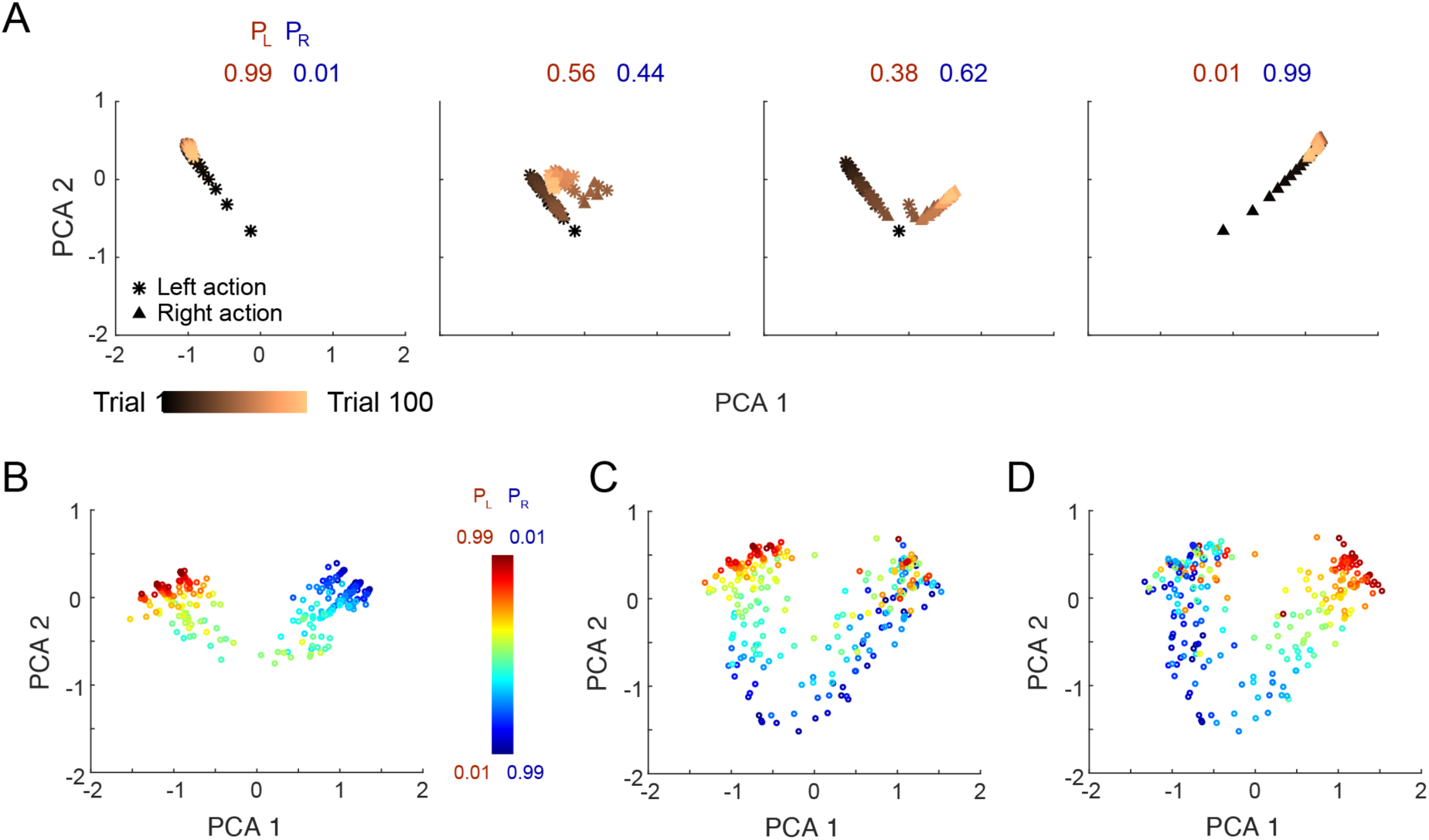
Visualization of RNN activations in “Illustrative example” (Bandit task). **A**) Evolution of RNN activation pattern during individual trials while testing on correlated bandits, after training with independent arm parameters. Format analogous to Figure 1E in the main text. *P_L_* = probability of reward for action *left*. *P_R_* = probability of reward for action *right*. Scatter points depict the first two principal components of the RNN activation (LSTM output) vector across steps of the bandit task, drawn from single trials across a range of randomly sampled payoff parameter settings. As evidence accrues, the activation pattern follows a coherent path in one of two directions, corresponding to the arms of the bandit. Interestingly, in more difficult discrimination problems (smaller gap between two arm reward probabilities; middle two columns, also see Figure 1E), activity sometimes appears to initially move toward one extreme of the manifold in early trials, then later reverses when late-coming observations contradict initial ones, to end up at other extremity by the end of the episode (see Ito and Doya, *PLoS Comput Biol, 11*, e1004540, 2015), which posits a related mechanism, though without t discussing its origins). **B**) RNN activity patterns from step 100 (episode completion) while testing on correlated bandits, across a range of payoff parameters. Format analogous to Figure 1F. In contrast to Figure 1F, in which the network is trained on linked arm parameters and thus the activity pattern traces out a one-dimensional manifold, the activity pattern after having trained on independent bandits is correspondingly more complex. **C**) RNN activity patterns when testing on independent bandits using the same network as in A and B, colored according to the left arm probability of reward (red = higher). **D**) same data as C, but colored according to the right arm probability of reward PR (red = higher).

**Supplementary Figure 2.**
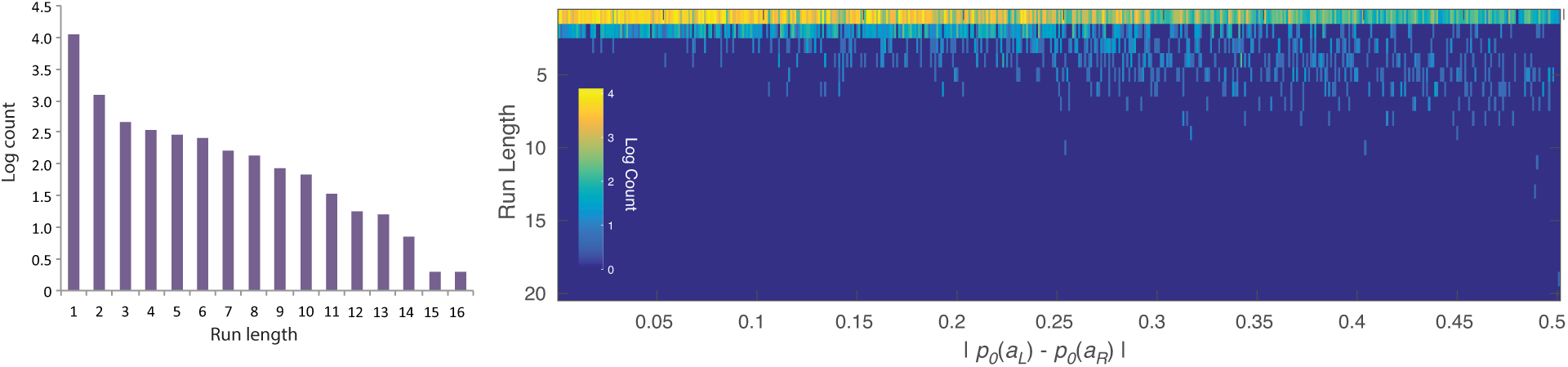
Detailed behavioral analysis of Simulation 1: Reinforcement learning in the prefrontal network. **Left:** Run lengths from the model. Bars indicate log trial counts for each run length, based on data from the last two-thirds of trials in each episode, and pooling across all reward-probability assignments. The pattern fits with the exponential decay typically reported in matching experiments (Corrado, Sugrue, Seung and Newsome, *Journal of the experimental analysis of behavior, 84*, 581–617, 2005), paired with a tendency to alternate responses. The latter is typical in tasks not involving a changeover penalty, and was observed in the study by Tsutsui et al. (Fabian Grabenhorst, personal communication). **Right:** Histograms showing log counts across run lengths individually for a range of task parameterizations. *P_0_ (a_i_)* denotes the ground-truth reward probability for action *i*. The pattern appears bimodal toward the right of the figure, suggesting that the model may approximate a fixed alternation between poorer and richer arms, remaining at the poorer arm for only one step and at the richer arm for a fixed number of steps. Houston and McNamara (Houston and McNamara, *Journal of the experimental analysis of behavior, 35*, 367–396, 1981) have shown that the optimal policy for discrete-trial variable-interval tasks, like the one studied here, takes this form.

**Supplementary Figure 3.**
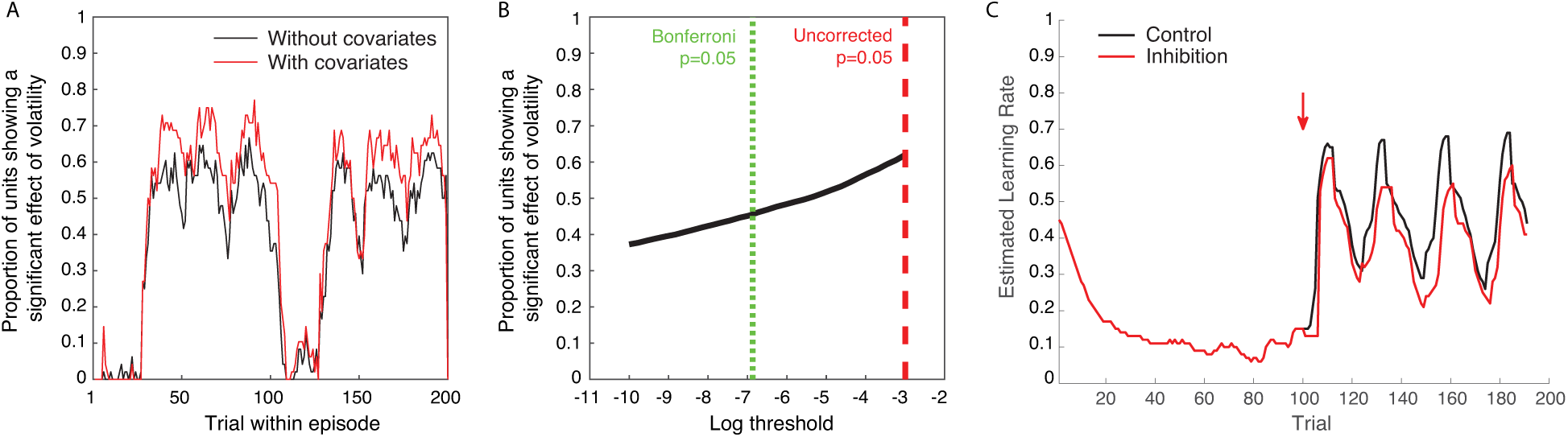
Additional simulations and analyses for Simulation 2: Adaptation of prefrontal-based learning to the task environment. **A**) Proportion of LSTM units coding for volatility changes over the course of an episode. Y-axis shows the fraction of units whose activity significantly correlated (across episodes) with the true volatility. Volatility coding emerged shortly after the 25th trial, which is when the first reversal occurred if the episode began with high volatility. Volatility coding was assessed by regressing hidden activations against the true volatility in a single regression (black line), or by multiple regression of hidden activations against true volatility, previous reward, next action, and true reward probability (red line). A unit was labelled as coding volatility if the slope of the regression was significantly different from zero at a threshold of 0.05/n, where n=48 is the number of hidden units. **B**) Because units were not independent, the true correction for multiple comparisons lies somewhere between 1 and 1/n. The significance threshold, for declaring a unit as coding volatility, varies on the x-axis in this plot, with uncorrected and Bonferroni corrected levels indicated by dashed lines. The y-axis shows the mean over trials of number of units that exceeded this threshold. Although changing the threshold had some effect on the number of units significantly coding volatility, there was a population of units that appeared to not code volatility at all, and another population that coded it very strongly. **C**) Causal role of volatility-coding units in controlling learning rate. After identifying the two units whose activity correlated most strongly (positively) with volatility, we inhibited these units by deducting a fixed quantity (0.3) from their activity level (LSTM cell output), analogous to an optogenetic inhibition. The black curve shows the learning rate (estimated as described in Methods) without this “optogenetic” manipulation. The red curve shows the learning rate when “optogenetic” inhibition was applied starting at the onset of the high volatility period (arrow). Inhibition of these two units resulted in a slower measured learning rate, establishing a causal connection between the volatility-coding units and learning dynamics. Note that we treated the relevant ‘pretraining’ to have occurred outside the lab, and to reflect statistics of naturalistic task situations. For additional related work on the neural basis of learning rate adjustment, see Soltani et al., *Neural Networks, 19*, 1075–1090, 2006; and Farashahi et al., *Neuron, 94*, 401–414, 2017. Like the comparable work cited in the main text, these studies posit special-purpose mechanisms. In contrast, our theory posits the relevant mechanisms as one manifestation of a more general meta-learning phenomenon.

**Supplementary Figure 4.**
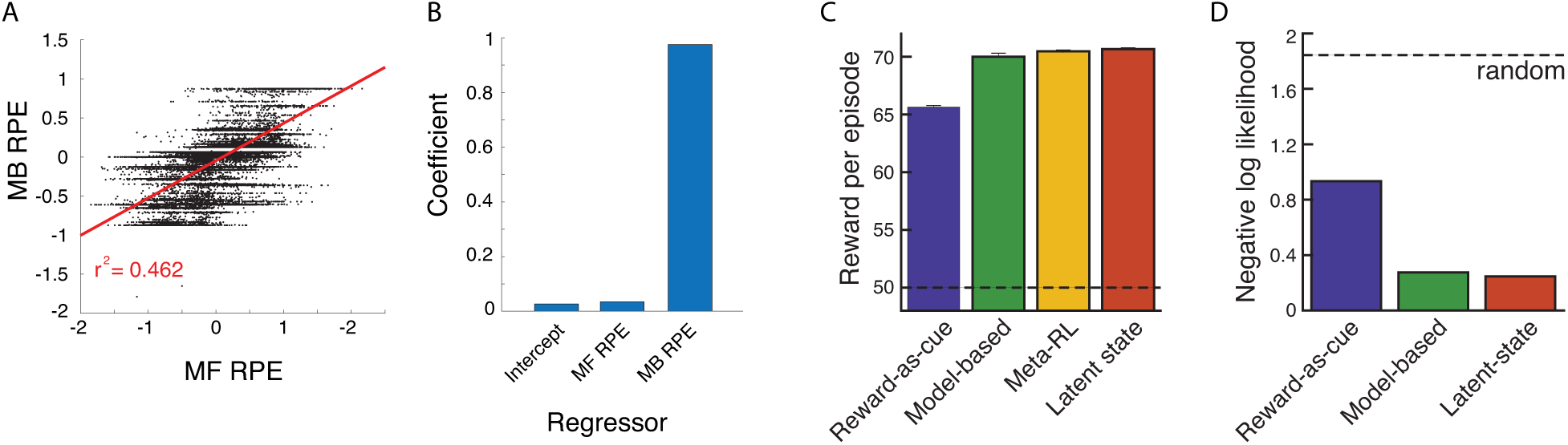
Additional analyses for Simulation 4: Model-based behavior: The Two-Step Task. **A**) Model-based and model-free RPEs, derived from prospective planning and SARSA, respectively, were partially correlated with one another. (This motivated the hierarchical regression in our primary analyses; see Figure 5F,G in main text.) **B**) Beta coefficients from multiple regression using both MF and MB RPEs to explain meta-RL’s RPEs. Only MB RPEs significantly explained meta-RL’s RPEs. **C**) Two-step task performance for meta-RL, model-based RL, and two alternative models considered by Akam et al. (*PLoS Comput Biol 11*, e1004648, 2015). Akam and colleagues (see also Miller et al., *bioRxiv 096339*, 2016) have shown that a reward-by-transition interaction is not uniquely diagnostic for model-basedness. It is possible to construct agents using model-free learning, which display a reward-by-transition interaction effect on the probability of repeating an action. Here we simulated and fit two models from Akam et al. (2015): (1) the “latent-state” model, which does inference on the binary latent state of which stimulus currently has high reward probability and learns separate Q-values for two the different states, and (2) the “reward-as-cue” model, which uses for its state representation the cross-product of which second-stage state occurred on the previous trial and whether or not reward was received. The model-based and latent-state strategies yielded comparable levels of reward, while the reward-as-cue strategy paid less well. Although the model-based agent earned slightly less than the latent-state agent, this difference was eliminated if the former was modified to take account of the anti-correlation between the second-step state payoff probabilities, as in the agent used to predict model-based RPE signals in Simulation 4 (see Methods). The performance of meta-RL was comparable to that of both the model-based and latent-state agents. We note that meta-RL is therefore unlikely to implement the reward-as-cue strategy; this is further ruled out by regression results presented in Figure 5E, which are inconsistent with the pattern produced by reward-as-cue (see Akam et al., 2015, and Miller et al., 2016). The dashed line indicates performance for random action selection. Model free Q-learning performed significantly worse than reward-as-cue (data not shown). **D**) Fitting three models to meta-RL’s behavior. Model-based and latent-state strategies yielded comparable fits, and again the small difference between them was eliminated if the model-based strategy was given access to a full model of the task. The reward-as-cue strategy fit significantly worse.

**Supplementary Figure 5.**
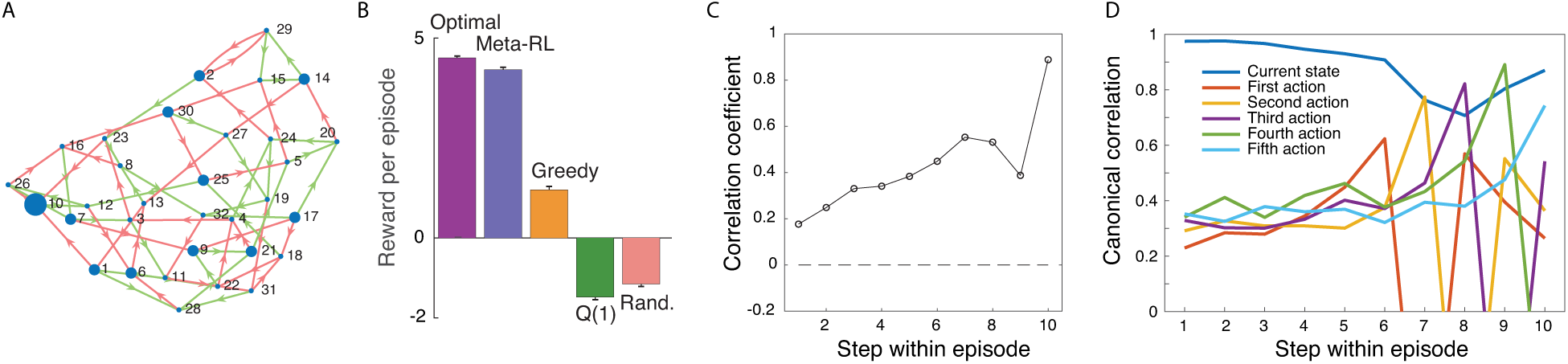
New simulation to directly test model-based reasoning in meta-RL. The two-step task alone is not sufficient to determine whether meta-RL can learn to implement truly prospective planning. Akam et al. (2015) have identified at least two non-prospective strategies (“reward-as-cue” and “latent-state”) that can masquerade as prospective in the two-step task. Although we can rule out meta-RL using the reward-as-cue strategy (see Supplemental Figure 4), we cannot rule out the latent-state strategy. (Akam and colleagues, 2015, acknowledge that the latent-state strategy requires a form of model to infer the current latent state from observed reward outcomes (Kolling et al., *Nature Neuroscience, 19*, 1280, 2016) – however, this is a relatively impoverished definition of model-based compared to fully prospective reasoning using a transition model.) However, other studies hint that true prospective planning may arise when recurrent neural networks are trained by RL in a multi-task environment. Recurrent neural networks can be trained to select reward-maximizing actions when given inputs representing Markov decision problem (MDP) parameters (e.g., images of random mazes) (Tamar et al., *Neural Information Processing Systems*, 2146, 2016; Werbos et al., *IEEE International Conference on Systems, Man and Cybernetics 1764*, 1996). A recent study by Duan and colleagues (*arXiv, 1611.02779*, 2016) also showed that recurrent networks trained on a series of random MDPs quickly adapt to unseen MDPs, outperforming standard model-free RL algorithms. And finally, recurrent networks can learn through RL to perform tree-search-like operations supporting action selection (Silver et al., *arXiv 1612.08810*, 2016; Hamrick et al., *Neural Information Processing Systems*, 2016; Graves et al., *Nature 538*, 471, 2016). We therefore sought to directly test prospective planning in meta-RL. To this end, we designed a novel revaluation task in which the agent was given five steps to act in a 32-state MDP whose transition structure was fixed across training and testing (for experiments using related tasks, see Kurth-Nelson et al., *Neuron, 91*, 194, 2016; Huys et al., *PNAS, 112*, 3098, 2015; Keramati et al., *PNAS, 113*, 12868, 2016; Lee et al., *Neuron, 81*, 687, 2014). The reward function was randomly permuted from episode to episode, and given as part of the input to the agent. Because planning requires iterative computations, the agent was given a ‘pondering’ (Graves, *arXiv:1603.08983*, 2016) period of five steps at the start of each episode. **A**) Transition structure of revaluation task. From each state, two actions were available (red and green arrows). For visualization, nodes are placed to minimize distances between connected notes. Rewards were sampled on each episode by randomly permuting a 32 element vector containing ten entries of +1, 21 entries of −1, and one entry of +5 – node size in diagram shows a single sampled reward function. There were over a billion possible distinct reward functions: orders of magnitude more than the number of training episodes. Episodes began in a random state. Meta-RL’s network architecture was identical to our other simulations, except the LSTM had 128 units. **B**) Meta-RL reached near optimal performance, when tested on reward functions not included in training. Significantly, meta-RL outperformed a ‘greedy’ agent, which followed the shortest path toward the largest reward. Meta-RL also outperformed an agent using the successor representation (data not shown). Q(1): Q-learning with eligibility trace; Rand.: random action selection. Bars indicate standard error. **C**) Correlation between network value output (‘baseline’) and ground-truth future reward grows during pondering period (steps 1 through 5). This indicates that the network used the pondering period to perform calculations that steadily improved its accuracy in predicting future reward. **D**) Canonical correlation between LSTM hidden state and several task variables, performed independently at each time step. Because “last action” was given as part of the input to the network, we orthogonalized the hidden state against this variable before calculating the canonical correlation. Unsurprisingly, we found that a linear code for action appeared most robustly at the step when the action was taken (the first action occurred on step 6, after five steps of ‘pondering’). However, this signal also ramped up prior to the onset of action. (The network cannot have a perfect representation of which action it will take, because the network’s output is passed through a noisy softmax function to determine the actual action given to the environment.) The network also maintained a strong representation of which state it currently occupied. We note that the network’s knowledge of the transition probabilities was acquired through RL. However, our theory does not exclude the existence of other neural learning mechanisms capable of identifying transition probabilities and other aspects of task structure. Indeed, there is overwhelming evidence that the brain identifies sequential and causal structure independent of reward (Glascher et al. *Neuron, 66*, 585, 2010; Tolman et al., *Psychological Review, 55*, 189, 1948). In subsequent work, it will be interesting to consider how such learning mechanisms might interact with and synergize with meta-RL (e.g., Hamrick et al., *Neural Information Processing Systems*, 2016).

**Supplementary Figure 6.**
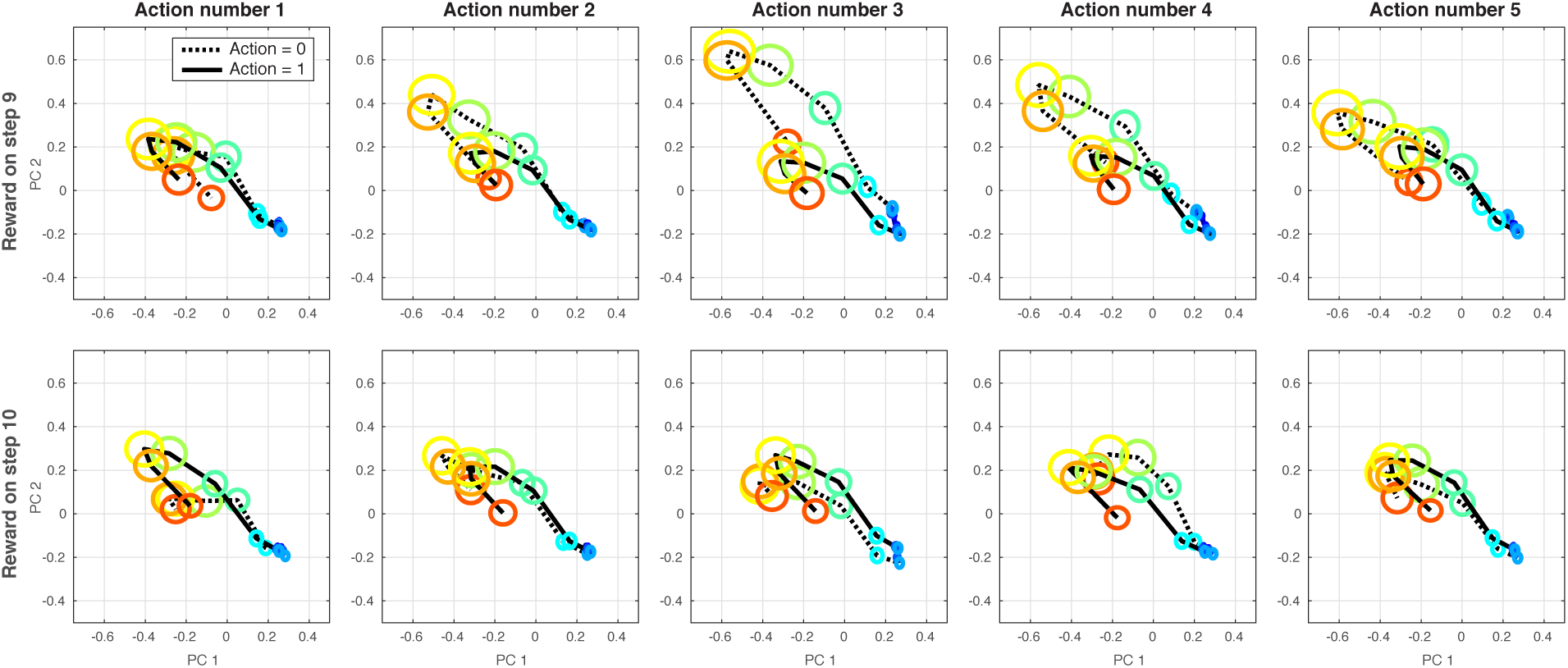
Trajectory of hidden activations during revaluation task (see Supplementary Figure 5). Each panel shows the evolution, over steps within an episode, of the first two principal components of the hidden activations of the LSTM network. Each oval is one time step. The width and height of each oval illustrates the standard error (across episodes), and the color corresponds to the index of the time step within the episode (blue at the beginning of the episode and red at the end). Each panel summarizes a subset of episodes. Episodes were randomly generated under the constraint that the large (+5) reward could be reached within exactly four or five steps from the initial state. The top row of panels shows all the episodes where the large reward was reached on the ninth time step (five ‘pondering’ steps followed by four overt actions). The bottom row shows the episodes where the large reward was reached on the tenth time step (pondering plus five actions). Together these comprise 97% of all episodes. The remaining 3% of trials, where the large reward was not reached, are not depicted. Within each row, the *i*-th panel (from left to right) divides the depicted episodes into two groups: those in which the *i-*th action ultimately executed was 0 (dotted line), and those in which it was 1 (solid line). We found strong interactions between the future reward and the current and future actions. For example, the network’s trajectory along the 2nd principal component began to diverge during the pondering period depending on whether it would take action 0 or 1 on the 3rd action. By the final step of pondering, this effect itself was strongly modulated according to whether reward would be received on step nine or ten of the episode. This appears to reflect a sophisticated but idiosyncratic planning algorithm.

**Supplementary Figure 7.**
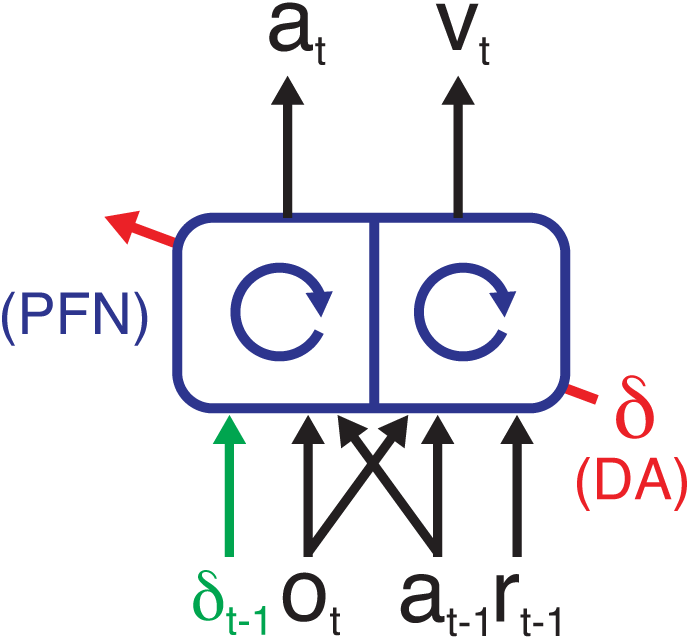
Agent architecture for Simulation 6. Agent architecture employed in Simulation 6. Instead of a single LSTM, we now employ 2 LSTMs to model an actor (outputting the policy) and a critic (outputting the value estimate). The critic receives the same input as in our original model, while the actor receives the state observation, last action taken, and the reward prediction error that is computed based on the value estimate output from the critic.

**Supplementary Figure 8.**
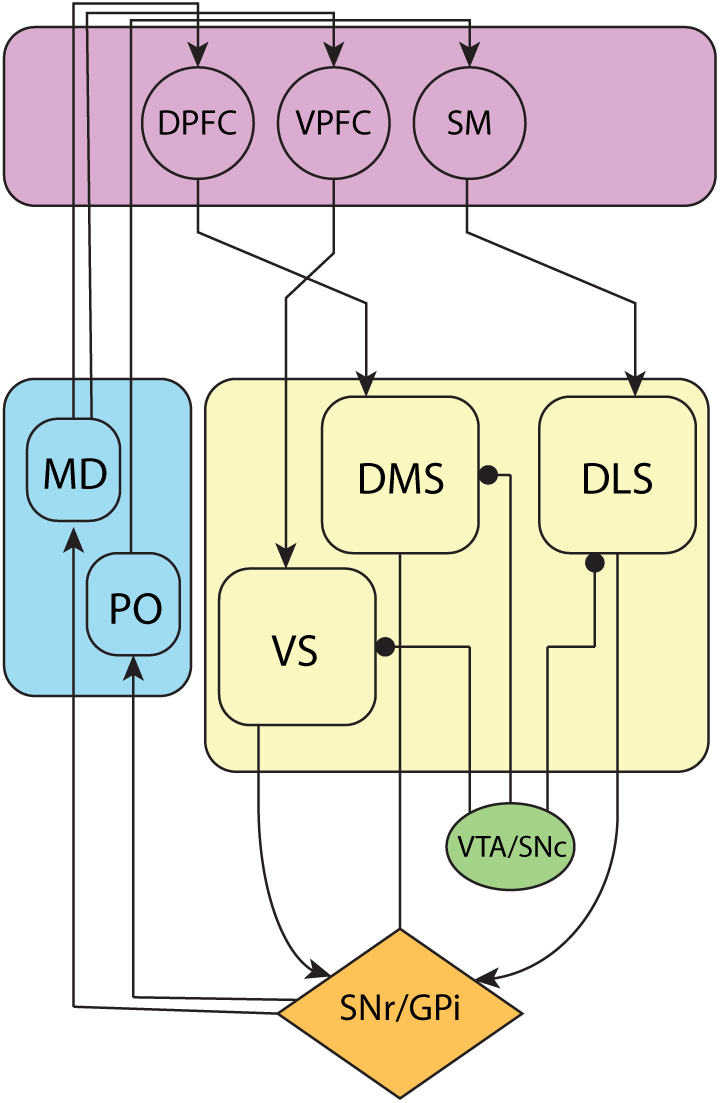
Cortico-basal ganglia-thalamic loops. As noted in the main paper, the recurrent neural system to which we refer as the prefrontal network includes not only the PFC itself, including orbitofrontal, ventromedial frontal, cingulate, dorsolateral prefrontal and frontopolar regions, but also the subcortical structures to which it most strongly connects, including ventral striatum, dorsomedial striatum (or its primate homologue), and mediodorsal thalamus. The recurrent connectivity that we assume is thus not only inherent within the prefrontal cortex itself but also spans parallel cortico-basal ganglia-thalamic loops. The figure diagrams a standard model of these loops, based on Glimcher, P. W., & Fehr, E. (Eds.). (2013). *Neuroeconomics: Decision making and the brain*. Academic Press, page 378. *DPFC*: Dorsal prefrontal cortex, including dorsolateral prefrontal and cingulate cortices. *VPFC*: Ventral prefrontal cortex, including ventromedial and orbitofrontal prefrontal cortices. *SM*: (Somato-) sensorimotor cortex. *VS*: Ventral striatum. *DMS*: Dorsomedial striatum. *DLS*: Dorsolateral striatum. *VTA*/*SNc*: Ventral tegmental area and substantia nigra pars compacta, with rounded arrowheads indicating dopaminergic projections. *SNr/GPi*: Substantia nigra pars reticulate/Globus pallidus pars interna. *MD*: Mediodorsal thalamus. *PO*: Posterior thalamus. The circuit running through DPFC has been referred to as the “associative loop”; the circuit running through the VPFC as the “limbic loop”; and the circuit running through SM as the “sensorimotor loop” (Haber, S., *Neuroscience, 282*, 248–257, 2014). One way of organizing the functions of the various regions of PFC and associated sectors of the striatum is by reference to the actor-critic architecture introduced in RL research, with ventral regions performing the critic role (computing estimates of the value function) and dorsal regions performing the role of the actor (implementing the policy; see Botvinick, M. M., Niv, Y. & Barto, A. C., *Cognition, 113*, 262–280, 2009, and Joel, D., Niv, Y. & Ruppin, E., *Neural Networks, 15*, 535–547, 2002). It is no coincidence that our assumptions concerning the outputs of the prefrontal network include both value estimates and actions. In this sense, we are conceptualizing the prefrontal network in terms of the actor-critic schema, and further elaborations of the paradigm might attain finer-grained architectural differentiation by importing existing ideas about the mapping between neuroanatomy and the actor-critic architecture (see, e.g., Song, H. F., Yang, G. R. & Wang, X.-J., *eLife, 6*, e21492, 2017). The implementation that we introduced in Simulation 6 takes a step in this direction, by dividing the prefrontal network into two halves, corresponding functionally to actor and critic. The loss function employed in optimizing our networks includes terms for value regression and policy gradient. Actor-critic models suggest that something very much like these two losses is implemented in cortex and basal ganglia (e.g. Joel, Niv & Ruppin, *Neural Networks, 15*, 535–547, 2002). Value regression is believed to be implemented in the basal ganglia as dopamine prediction errors drive value estimates toward their targets (Montague, Dayan & Sejnowski, *Journal of Neuroscience, 16*, 1936–1947, 1996). Meanwhile, the policy gradient loss simply increases the strength of connections that participated in generating an action (when that action led to positive reward), which is believed to be the role of dopamine acting as modulator of plasticity at corticostriatal synapses. Under the meta-RL theory, the role of DA in modulating synaptic function should play out only over a relatively long time-scale, serving to sculpt prefrontal dynamics over the course of multiple tasks. This is in contrast to the standard model, which assumes that DA can drive synaptic change sufficiently quickly to affect behavior on the scale of a few seconds. Interestingly, despite decades of intensive research, no direct evidence has yet been reported to indicate that phasic DA signals can drive synaptic change this rapidly. Indeed, we are aware of no experimental evidence that phasic elevations in DA concentrations spanning less than a full minute can significantly impact synaptic efficacy, and in many studies the impact of phasic fluctuations in DA can take minutes (or even tens of minutes) to ramp up (Brzosko, Z., Schultz, W. & Paulsen, O., *Elife, 4*, e09685, 2015; Otmakhova, N. A. & Lisman, J. E., *Journal of Neuroscience, 16*, 7478–7486, 1996; Yagishita, S. et al., *Science, 345*, 1616–1620, 2014). The lack of evidence for faster effects of phasic DA on synaptic plasticity presents another difficulty for the standard model. In contrast, the meta-RL framework directly predicts that such short-term effects of DA should in fact be absent in the prefrontal network, and that the effect of phasic DA signaling on synaptic efficacy in this network should instead operate on a significantly longer time-scale.

**Supplementary Video 1.** Video illustrating performance in Simulation 5: Learning to learn (https://youtu.be/FXxAobxqPUo).

Four consecutive trial-blocks are shown. In the first and second blocks the rewarded image first appears on the left; in the third and fourth on the right. Images are shown in full resolution. See Methods for details concerning simulation methods.

